# MicroRNA regulation of the MRN complex impacts DNA damage, cellular senescence and angiogenic signaling

**DOI:** 10.1101/132258

**Authors:** Cristina Espinosa-Diez, RaeAnna Wilson, Namita Chatterjee, Clayton Hudson, Rebecca Ruhl, Christina Hipfinger, Erin Helms, Omar F. Khan, Daniel G. Anderson, Sudarshan Anand

## Abstract

MicroRNAs contribute to biological robustness by buffering cellular processes from external perturbations. Here we report an unexpected link between DNA damage response and angiogenic signaling that is buffered by two distinct microRNAs. We demonstrate that genotoxic stress-induced miR-494 and miR-99b inhibit the DNA repair machinery by targeting the MRE11a-RAD50-NBN (MRN) complex. Functionally, gain and loss of function experiments show that miR-494 and miR-99b affect telomerase activity, activate p21 and Rb pathways and diminish angiogenic sprouting *in vitro* and *in vivo*. Genetic and pharmacological disruption of VEGFR-2 signaling and the MRN complex reveal a surprising co-dependency of these pathways in regulating endothelial senescence and proliferation. Vascular-targeted delivery of miR-494 decreases both growth factor-induced and tumor angiogenesis in mouse models. Mechanistically, disruption of the MRN complex induced CD44, a known driver of senescence and regulator of VEGF signaling in addition to suppressing IL-13 a stimulator of VEGF signaling. Our work identifies a putative miR-facilitated mechanism by which endothelial cells can be insulated against VEGF signaling to facilitate the onset of senescence and highlight the potential of targeting DNA repair to disrupt pathological angiogenesis.

## Introduction

Accumulation of DNA damage can overwhelm the repair machinery and lead to senescence (1). Endothelial senescence leads to progressive damage and deterioration of cellular structure and function over time (2) (3). Two major pathways of senescence in endothelial cells are replicative senescence and stress induced premature senescence (SIPS). Replicative senescence is one of the hallmarks of aging and is associated to telomere shortening. Stress-induced premature senescence is triggered by external stimuli, including oxidizing agents and radiation, both of which can induce DNA damage and cell cycle arrest.

Recent studies indicate that DNA repair proteins in endothelial cells (ECs), such as ATM kinase and histone H2AX, have an inherently pro-angiogenic role (4, 5). For example, ATM deficiency kinase decreases tumor angiogenesis while enhancing the anti-angiogenic action of VEGF blockade, suggesting that pathological neoangiogenesis requires ATM. ATM can also function as a redox sensor independent of its DDR function and regulate oxidative stress responses (6) and was recently implicated as a driver of cellular senescence (7). Similarly, global or EC specific deletion of the histone H2AX in mice results in a substantial decrease of pathological angiogenesis in proliferative retinopathy, hindlimb ischemia and tumor angiogenesis. These findings suggest that key regulators of DNA damage repair (DDR) modulate pathological angiogenesis.

Mre11a-RAD50-NBN (MRN) complex acts as a sensor of DNA double strand breaks (DSB), initiating Homologous Recombination (HR) or Non-Homologous-End-Joining (NHEJ) pathway. Once MRN detects DSBs, it tethers the ends, activates and recruits DNA damage response proteins such as ATM (8) (9). Interestingly, MRN is also associated with telomere maintenance, playing a role in the formation and disassociation of the t-loops (10, 11). The MRN complex relationship with aging and cell senescence has been described (12, 13), however its role in ECs and pathological neovascularization is unclear. Similarly, while several miRs have been shown to be involved in EC senescence (2, 14, 15), proliferation, viability and migration (16) (17) (18), our work identifies a novel link between DDR driven induction of miR-494, miR-99b, the MRN complex, EC senescence and tumor angiogenesis. Our results indicate that altering the DDR response in tumor ECs, not only enhances cellular senescence but also has a functional effect on both growth factor induced and pathological angiogenesis leading to decreased tumor growth..

## Results

### DNA damage induces miRs-494 and miR-99b *in vitro* and *in vivo*

We previously reported a seven microRNA signature (miR-103, miR-494, miR-99b, miR-21, miR-224, miR-92a and let-7a) specifically upregulated in ECs after radiation, hydrogen peroxide and cisplatin treatment. Among these, we found that miR-494 and miR-99b are both transcribed rapidly in human umbilical vein endothelial cells (HUVECs) in response to γ-radiation with maximal induction occurring at a lower (2 Gy) dose of radiation. (Figure 1A-B). Consequently, we observed that miR-494 was also upregulated after radiation in a PyMT breast cancer model both at low and high doses at different time points (Figure 1C-D).

**Figure 1:**
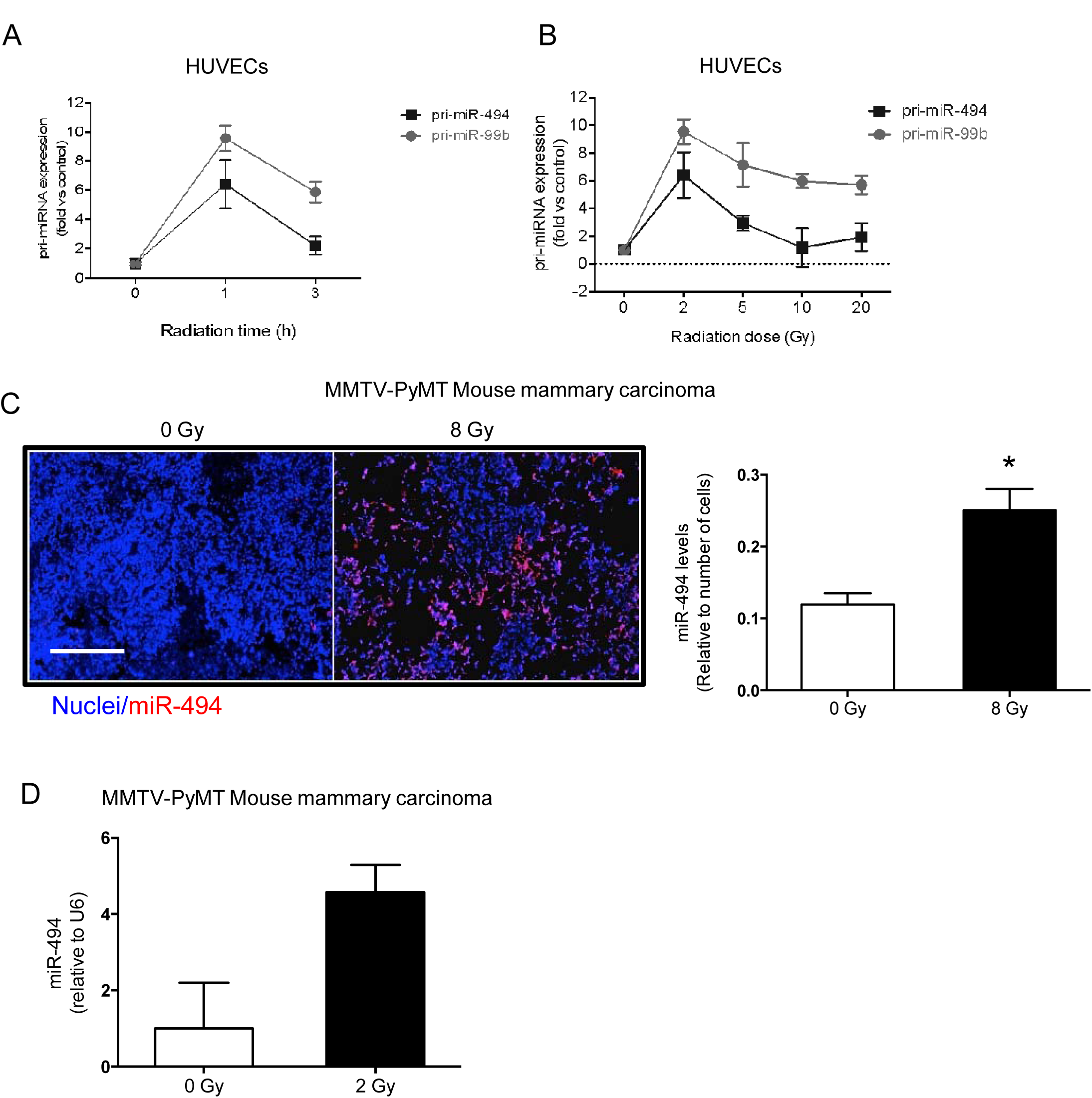
miR-494 and miR-99b are induced in response to radiation. **A)** Kinetics and **B)** Dose response of primary miR-494 and miR-99b transcripts in HUVECs treated with the indicated dose of radiation. The dose response was evaluated at 1h post radiation. **C)** In situ hybridization of miR-494 on FFPE slides from PyMT-MMTV tumors irradiated with the indicated dose of radiation. Tumors were harvested 36h post radiation. Bars depict mean + SEM of sections from 3 mice per group. * *P*<0.05; two-tailed Student’s T-test. **D)** qRT-PCR of mature miR-494 from orthotopic PyMT tumor implants (n= 4 tumors per group) at 6h post radiation. Bars depict mean + SEM.

### Gain and loss of miRs-494 and 99b affect pathological and replicative senescence

To understand the role of these two miRs, we performed different gain and loss of function assays in our endothelial cell model, HUVECs. We observed that the most prominent phenotype with these two miRs was the induction of cellular senescence. Gain of either miR-99b or miR-494 in early passage HUVECs increased senescence associated β-galactosidase levels (SA-β gal). This phenotype was also observed in human microvascular endothelial cells (HMVECs) and normal human lung fibroblasts suggesting this phenotype was not exclusive to venous endothelium (Supplementary Figure 1). In contrast, inhibition of miR-99b or miR-494 decreased SA-β gal levels significantly in HUVECs undergoing pathological senescence induced by a high-dose of radiation (Figure 2B). We asked if these miR inhibitors had an effect on replicative senescence and found that inhibition of either miR-99b or miR-494 in late passage senescent HUVECs decreased their β-gal and increased activation of caspase 3 and 7 (Figures 2C-D). Similarly, gain of miR-494 and miR-99b affected cell cycle progression whereas the inhibition of the miRs decreased the G2 arrest in response to a 10 Gy dose of radiation (Supplementary Figure 2). Interestingly, miR-494 expression decreased BrdU incorporation whereas miR-99b expression increased activation of Caspase-3 & 7 (Supplementary Figure 3). Given the strong influence of DNA damage on cellular senescence, we asked if either of these miRs affected DNA damage. Indeed, ectopic expression of both miR-494 and miR-99b increased histone H2AX phosphorylation, a marker of DNA damage (Figure 2E). Importantly, this increase in DNA damage resulted due to the miRs alone without any additional source of DNA damage. Consistent with these findings, we observed some critical features of senescence - a decrease in telomerase activity (Figure 2G), an increase in the cell cycle regulator p21 as well as a decrease in Rb hyperphosphorylation (Figure 2H). Moreover, Phalloidin staining in HUVECs after 48 hours of miR-494 treatment alsorevealed flattened and multinucleated cells (Figure 2F), another morphological phenotype associated with stress dependent senescence. Taken together, these observations establish that miR-494 and 99b drive EC senescence likely via pathways involved in DNA damage or repair.

**Figure 2.**
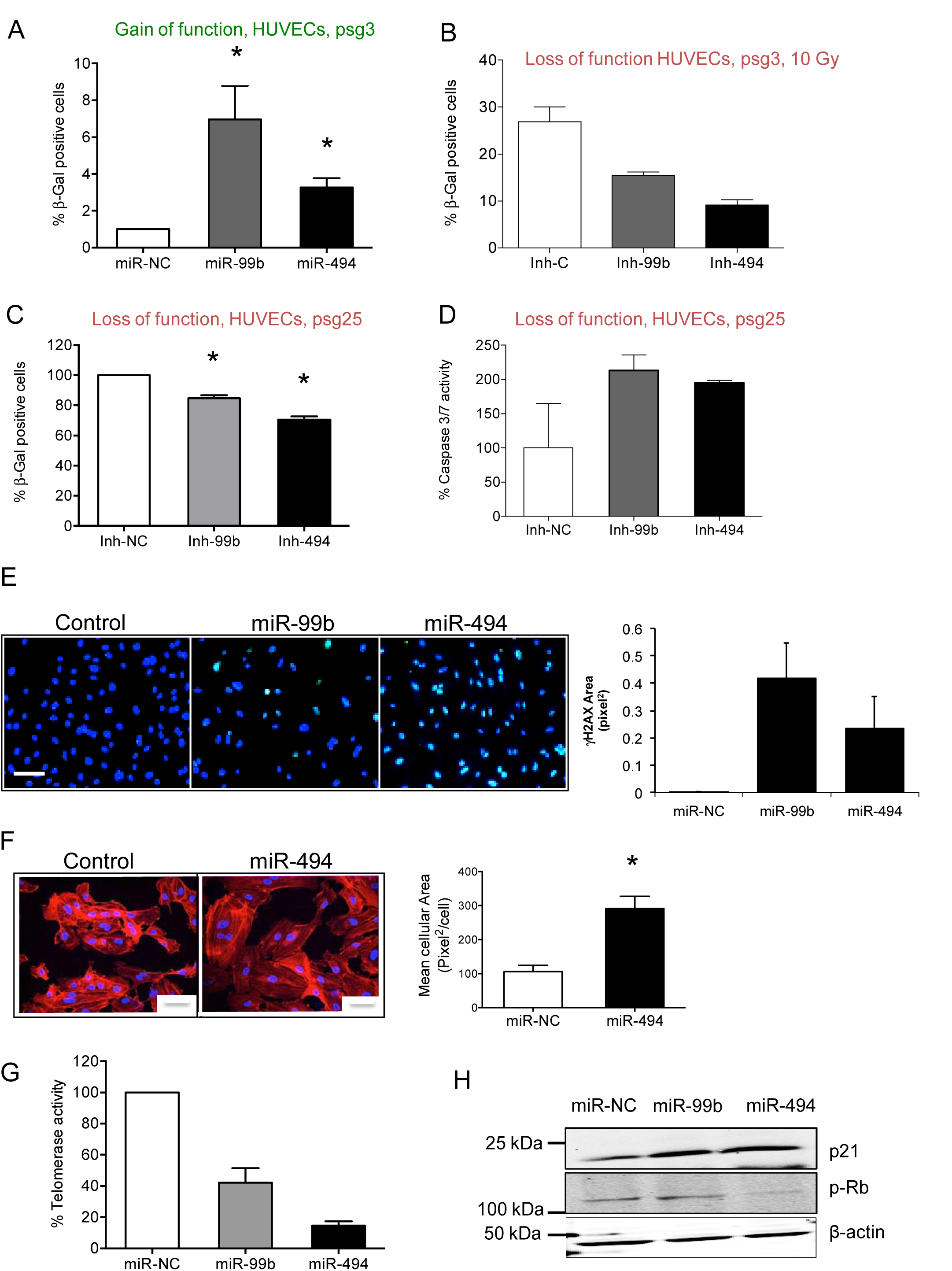
miR-494 and miR-99b drive pathological and physiological endothelial senescence by disrupting DNA repair. **A)** SA*β*-Gal assay in early passage HUVECs transfected with a miR negative control or miR-99b or miR-494 mimics. Bars show % (mean + SEM) of *β*-gal positive cells for at least hundred cells analyzed 48h post transfection. **B)** SA*β*-Gal assay at 48h post radiation of early passage HUVECs transfected with either an inhibitor control or inhibitors of miR-99b or miR-494. Bars show % (mean + SEM) of *β*-gal positive cells for at least hundred cells analyzed. **C-D)** SA*β*-Gal assay **(C)** and Caspase-3/7 activity **(D)** in senescent (psg 25) HUVECs transfected with either an inhibitor control or inhibitors of miR-99b or miR-494 at 48h post transfection. Bars show mean + SEM. **E)** γH2AX staining in HUVECs 48h after transfection with a miR negative control or miR-99b or miR-494 mimics. Bars depict mean γH2AX area per nuclear area + SEM of 3-4 technical replicates per group. **F)** HUVECs were transfected with either a miR negative control or miR-494 mimic. After 48h, cells were fixed and stained with anti-phalloidin antibody (red) and DAPI (blue) as indicated. Bars depict % mean cellular area + SEM. **G)** HUVECs were transfected with a miR negative control or miR-99b or miR-494 mimics. 48h later telomerase activity was assayed using a Telo-Tagg assay. Bars depict % mean + SEM. **H)** Representative western-blot of senescence marker p21 and pRB in HUVECs transfected as in G).

### miR-494 and miR-99b target the MRN DNA repair complex affecting senescence

To analyze the possible targets of miR-494 and miR-99b responsible for this phenotype, we ectopically expressed these two miRs in HUVECs and analyzed gene expression using a DNA damage array (Supplementary Figure 4). Interestingly, we identified three common targets: MRE11a, RAD50 and NBN. RNA hybrid modeling suggested putative binding sites for both miRs on all three target 3’UTR regions (Supplementary Figures 5-6). Both miR-494 and −99b directly bound MRE11A mRNA, and to a lesser extent, RAD50 and NBN mRNAs, as measured by a miRTrap assay (Clontech) (Figure 3A). Consistent with this finding, transfection of miR mimics decreased MRN RNA levels by qRT-PCR (Figure 3B) and protein levels as measured by a western blot (Figure 3C). We corroborated the interaction of these miRs with their targets by cloning the 3’-UTRs in luciferase reporter vectors, and co-transfecting them in HEK-293T cells. Either of the miRs transfected alone was able to decrease the luciferase activity downstream of the MRN 3’-UTR reporter (Figure 3D). Finally, miR-494 transfected HUVECs showed a decrease in MRE11a nuclear foci number (Figure 3E).

**Figure 3.**
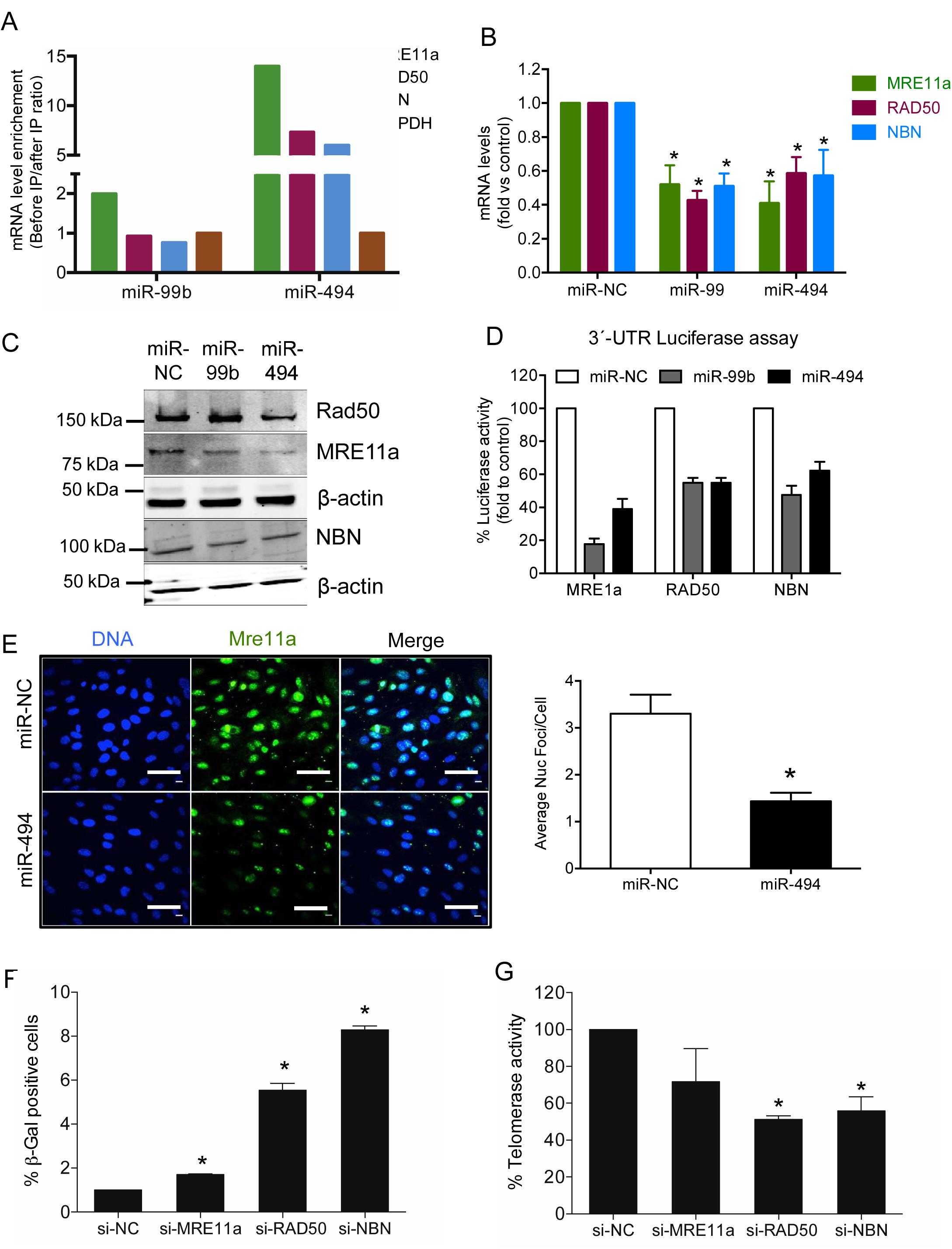
miR-494 and miR-99b target the MRN complex in ECs. **A)** miR-TRAP immunoprecipitation. Bars depict mean + SEM of mRNAs bound to the RISC complex quantified by qRT-PCR in HEK-293T cells transfected with either miR-99b or miR-494 for 24h. **B)** qRT-PCR depicting target mRNA levels in HUVECs transfected with either miR-negative control or miR-99b or miR-494. Bars show mean + SEM of the three MRN complex members. **C)** Representative western-blot of MRE11a, RAD50 and NBN after 48h transfection of indicated miRs in HUVECs. **D)** Luminescence from 3’-UTR-luciferase constructs for MRE11a, RAD50 and NBN 24h after transfection with miR-99b or miR-494. Graph represents mean +SEM of 3 independent experiments. **E)** HUVECs were plated on glass coverslips and 24h later transfected with either miR-494 or miR negative control. After 48h cells were fixed and stained with MRE11a antibody. Scale bar represents 10 μm. Bar graph depicts average foci/ cell + SEM. **F)***β*-Gal assay and **G)** Telomerase activity in HUVECs transfected for 48h with specific siRNAs against MRE11a, RAD50 and NBN or an siRNA negative control. Bars depict % mean + SEM. \**P*<0.05; two-tailed Student’s T-test.

miRs typically regulate several target mRNA transcripts making their functional attribution challenging (19), (20-23). To address this, we used a target protector, a locked nucleic acid (LNA) stabilized oligonucleotide, that binds completely to a specific miR-binding site on a single target mRNA, to rescue it from the negative regulation of the miR. We confirmed that the MRE11A target protector, rescued MRE11A RNA levels even in the miR-494 transfected cells compared to a control target protector oligo (Supplementary Figure 7A). Functionally, the MRE11A target protector significantly restored the telomerase activity and to a lesser extent the senescence associated *β*-gal levels in miR-494 transfected cells (Supplementary Figure 7B-C). We then sought to determine if disruption of the MRN complex using siRNAs (Supplementary Figure 8) resulted in a similar phenotype. Indeed, silencing each of the components of the MRN complex induced senescence and impaired telomerase activity (Figure 3 F-G). Confirming these observations, we found that siMRN as well as Mirin-1, a small molecule inhibitor specific to MRE11a (24), also enhanced senescence in HMVECs. These data indicate that the miR-494 and 99b both downregulate the MRN machinery and thereby both exacerbate DNA damage and drive senescence.

### The MRN pathway cross-talks with angiogenic growth factors

While disruption of DNA damage response (DDR) in ECs can modulate both pathological and developmental angiogenesis via VEGF (4, 5), it is not clear if targeting MRN would also influence VEGF signaling. Our results show that combinatorial disruption of both VEGFR2 and MRE11a with Mirin-1 and Vandetanib, a small molecule inhibitor for VEGFR2 (Figure 4A), led to a decrease in proliferation of HUVECs with a synergistic interaction at the lowest dose (Chou-Talalay Combination Index <0.4). Conversely, disruption of the MRN complex with Mirin-1 affected bFGF mediated pERK signaling and to a lesser extent VEGF induced pERK in HUVECs (Figure 4B). Interestingly, we silenced VEGF-receptor 2 (VEGFR2) and tested whether disruption of the MRN complex induced senescence. Surprisingly, knockdown of VEGFR2 diminished the senescence promoting activities of both the siRNAs targeting the MRN and Mirin-1 (Fig 4C-D). These experiments argue that the disruption of MRN could inhibit angiogenic growth factor signaling. Indeed, siRNA targeting of Mre11a or NBN significantly decreased angiogenic sprouting over a four days in a 3D fibrin matrix.

**Figure 4.**
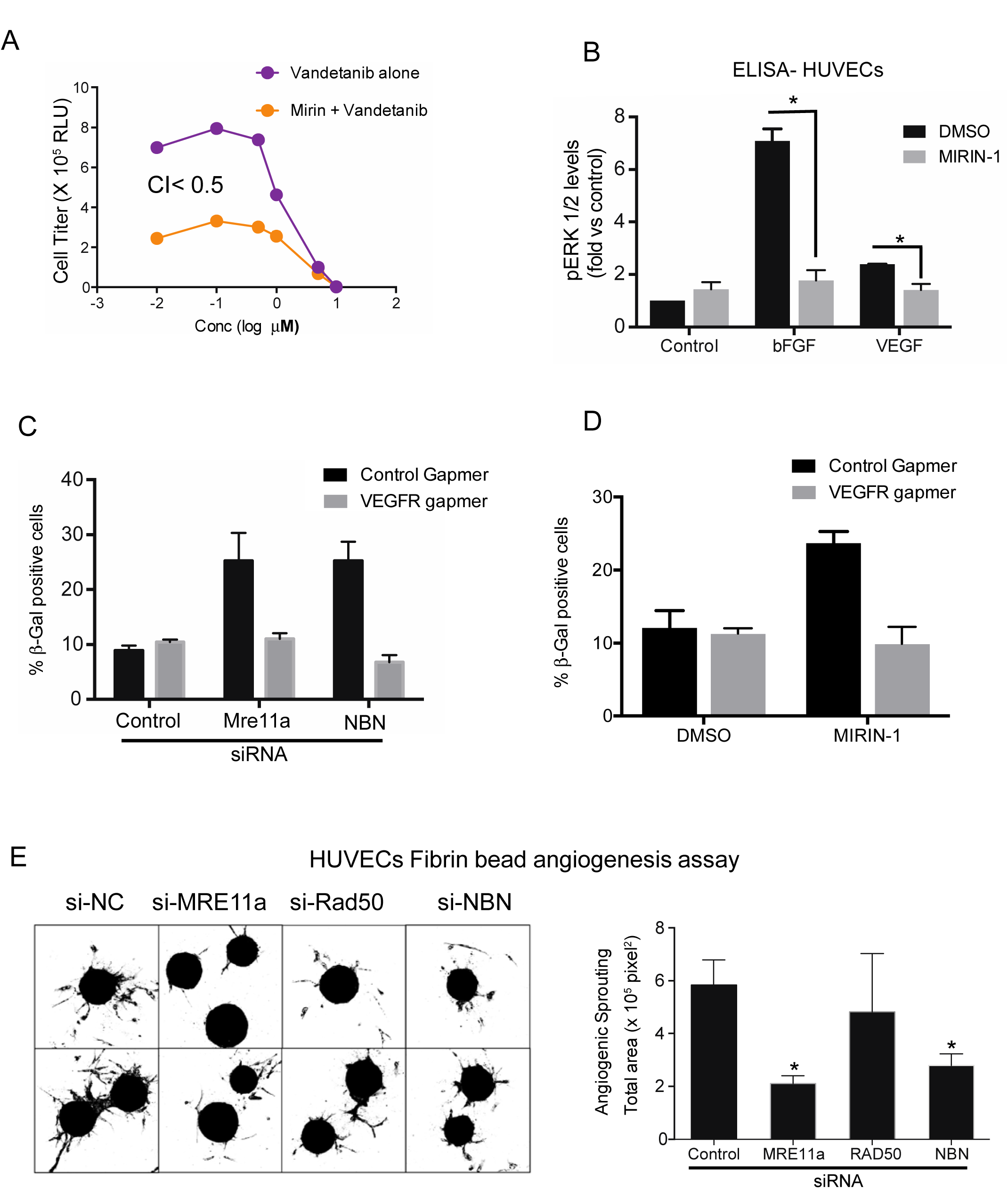
MRN pathway interacts with angiogenic signaling and is necessary for sprouting angiogenesis. **A**) Cell proliferation assay in HUVECs treated with VEGFR2 inhibitor Vandetanib (10 μM) alone or in combination with Mirin-1 (50 μM). Combination Index (CI) at the lowest concentration is shown. **B)** Representative pERK ELISA. HUVECs were starved overnight and treated with Mirin-1 (50 μM). 30 mins later cells were treated for 10 min with VEGF (50ng/ml) or bFGF (100ng/ml) and p-ERK levels were measured using an ELISA assay. Graphs depict mean p-ERK levels + SEM. **C) β**-Gal assay in HUVECs co-transfected for 48h with siRNAs against MRE11a or NBN in combination with a VEGFR2 silencing gapmer. Bars show % mean + SEM of β-gal positive cells for at least hundred cells analyzed. **D)** *β*-Gal assay in HUVEC transfected for 48h with VEGFR2 gapmer and treated with Mirin-1 (50μM) for 24h. Bars show % mean ± SEM of *β*-gal positive cells for more than 100 hundred cells analyzed. **E)** Fibrin bead angiogenesis assay. HUVECs were transfected with the indicated siRNAs and assessed for their sprouting angiogenesis potential. The images show representative lectin stained beads for each condition. Bars depict mean +SEM of lectin area analyzed across at least 25 beads per group. \**P*<0.05; two-tailed Student’s T-test.

### miRs-494 and 99b drive senescence through cell intrinsic and extrinsic mechanisms

To understand the shared molecular pathways that may drive senescence downstream of miR-494, miR-99b and the MRN complex, we utilized a human senescence gene signature array. We found 6 genes with similar changes across both the miR and siRNA treatment groups (Supplementary Fig 9A). Interestingly, CD44, a known negative regulator of VEGF signaling (25, 26), was the most significantly upregulated gene in the four groups. This increase in gene expression was also reflected by increased cell surface expression of CD44 (Supplementary Figure 9B). We also performed a cytokine expression profiling using a multiplex luminex assay and discovered that miR-494 robustly suppressed IL-13 levels. IL-13 can function as a driver of VEGF signaling (27) or via its shared IL-4 receptor leading to STAT6 phosphorylation and senescence (28). We are currently investigating the relative contributions of these pathways to miR-494 and MRN function in EC senescence.

### miR-494 mimic disrupts growth factor driven and pathological angiogenesis

Thus far, our data shows that miR-494 and 99b induce senescence by inhibiting the MRN complex, which diminishes VEGF/bFGF signaling, and sprouting angiogenesis. Next, we addressed whether this miR pathway can be modulated to regulate angiogenesis *in vivo*. First, we confirmed that the miR mimics miR-494 and 99b also disrupted sprouting angiogenesis in the fibrin bead assay (Figure 5A). Next, we used a growth factor induced angiogenesis model by implanting bFGF containing Matrigel plugs in mice. We treated the mice with either a miR-494 mimic or a control mimic in vascular-targeted 7C1 nanoparticles (29, 30). We found a robust decrease in CD31 staining in miR-494 treated plugs compared to the control miR mimic treated group (Figure 5B). To test the utility of this approach in a pathological angiogenesis, we utilized a highly aggressive, triple negative 4T1 mouse mammary carcinoma model. Treatment of vascularized tumors with miR-494 in the 7C1 nanoparticles resulted in a delayed tumor growth for a few days (Figure 5C). We also observed a trend towards a decrease in lung metastases (Figure 5D). Importantly, we observed the miR-494 treated tumors had a significant decrease in angiogenesis as measured by CD31 staining (Figure 5E). Our in vivo studies indicate that miR-494 is able to decrease both growth factor induced and pathological angiogenesis *in vivo*.

**Figure 5.**
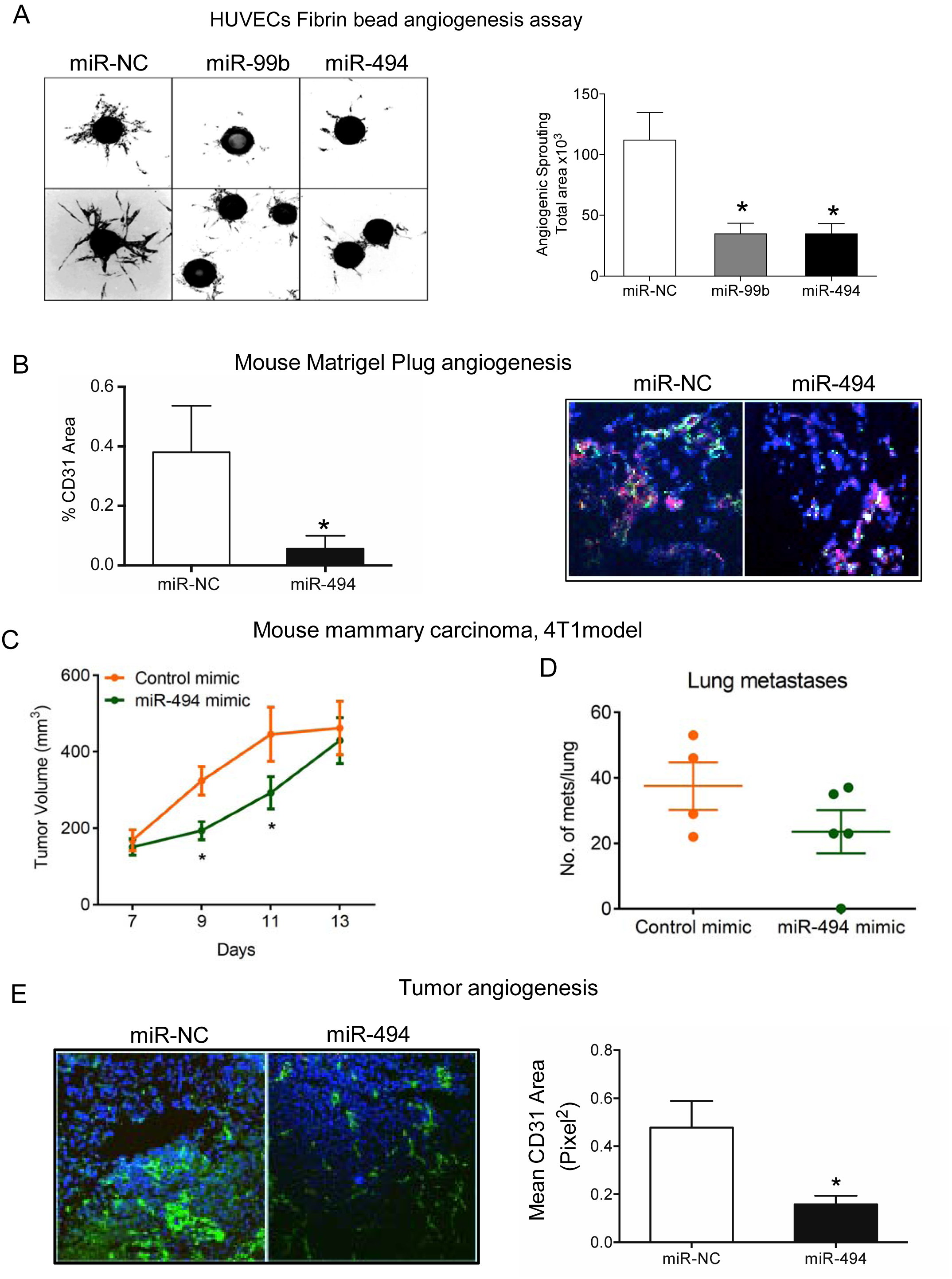
miR-494 disrupts angiogenesis *in vitro* and *in vivo*. **A)** Fibrin bead angiogenesis assay. HUVECs were transfected with the indicated miRs and assessed for their sprouting angiogenesis potential. The images show representative lectin stained beads for each condition. Bars depict mean +SEM of lectin area analyzed across at least 25 beads per group. **B)** CD31 staining *in vivo*. bFGF containing Matrigel plugs were implanted subcutaneously in nude mice and treated with miR-NC or miR-494 in vascular targeted 7C1 nanoparticles on days 5 and 6. Mice were sacrificed at day 7 and plugs were harvested for tissue sections. Angiogenesis was measured by staining sections with anti-CD31 (green) and αSMA (red) and DAPI (blue). Quantification of CD31 area from at least three mice per group is shown. Bars show mean + SEM. C) 4T1 mouse mammary carcinoma cells (1X10^4^) were implanted in the mammary fat pads of 6-8 week old female Balb/C mice (N= 4-5 per group). Mice were randomized to receive either Control miR mimic or miR-494 mimic formulated as miR-7C1 nanoparticles (0.7 mg/kg, i.v.) on days 12, 14, 16. Primary tumor volume measurements (C) and gross metastatic foci per lung on day 20 (D) are shown. Error bars depict SEM. * P<0.05 by Mann-Whitney U-test. E) Angiogenesis was measured by staining the 4T1 tumor sections with anti-CD31. Bars show CD31 area normalized the tumor area as mean + SEM from n=3 tumors per group.

These results demonstrate that the DNA repair pathway and the MRN complex have an essential functional role in angiogenesis and we propose that we can exploit this DNA repair-VEGF pathway codependency for future antiangiogenic therapies. Genotoxic stress agents induce DNA damage in ECs; however, they also increase expression of miR-494 and miR-99b. These two miRs lead to senescence by targeting the DNA repair complex MRN. Moreover, disruption of MRN decreases proliferation and ERK phosphorylation in response to angiogenic growth factors. In addition, other effects such as CD44 induction and IL-13 suppression can also amplify the induction of senescence and inhibition of angiogenesis.

## Discussion

We have postulated that a network of stress-dependent miRs target DNA repair pathways leading to impaired angiogenesis (29). Among this group of stress-induced miRs, miR-494 and miR-99b lead to senescence, likely through MRN disruption. miR-494 has been shown to be highly upregulated in retinoblastoma (31), cardiovascular pathologies such as cardiac injury (32) and in atherosclerotic lesion development (33). miR-494 has also been reported as an angiogenesis inhibitor, supporting our data here (34-36). Recently, Esser et al demonstrated that miR-494 is downregulated in ECs treated with the pro-angiogenic factor BMP4, in opposition to an increase in the pro-angiogenic miR-126 (37). miR-494 has been implicated in tumor senescence and the development of resistance to radiation and chemotherapy (38) (39) (40) (41).

In EC-progenitors, miR-99b affects the expression of PECAM-1 and VE-cadherin (42). miR-99b also modulates β-catenin in endothelial junctions (43). Interestingly, miR-99b expression has been proposed as a prognostic marker to assess the response to VEGFR2 Tyrosine Kinase Inhibitors (TKI). Lukamowicz-Rajska et al described higher expression of miR-99b-5p in clear cell renal cell carcinoma (CCRC) patients with complete response to TKI treatments (44) compared to partial or non-responders. Another example is dermal wound healing, where TK pathways have decreased signaling. Prior studies proposed that miR-99b targets several members of the IGFR1/AKT/mTOR signaling pathway. They demonstrated that miR-99b was downregulated in the early phases of wound healing likely facilitating cell proliferation (45). We propose here that miR-99b expression is leading to senescence through MRN disruption, but that its effect on the VEGFR2/ERK signaling pathway enhances the anti-angiogenic phenotype. Similarly, our observations of miR-99b diminishing a specific VEGF-dependent early-gene signature (46), also suggests that the induction of miR-99b in complete responders can enhance the effect of VEGFR2 TKIs.

We find that miR-494 and miR-99b target MRE11a, NBN and RAD50 to affect EC senescence and sprouting angiogenesis. EC senescence leads to dysfunction in cardiovascular diseases (47, 48). Loss of function mutations in the MRN genes cause inherited genetic disorders that are characterized by elevated sensitivity of patients’ cells to radiation damage (49, 50). Interestingly, there is some evidence that NBN disruption has an anti-angiogenic effect *in vivo* (51). Our observations also imply that the senescence phenotype maybe mediated by the increase in CD44 expression on the cell surface (52). CD44 is a known regulator of endothelial interactions with leukocytes in addition to its other critical roles in EC junction integrity (53). Induction of CD44 in response to DNA damage by miR disruption of the MRN complex may increase adhesiveness of the endothelium (54, 55). Similarly, there are demonstrated roles for IL-13 in enhancing VEGF signaling (27) or senescence (28) that are plausible downstream of miR-494 regulation of the MRN pathway.

Emerging studies have implicated deficient telomere maintenance as leading to senescence when MRN function is compromised. For example, decreased MRE11a has been shown to increase T-cell aging in arthritis due to compromised telomere maintenance and heterochromatin unraveling (56). Recent structural studies also shed light on how the phosphorylation of NBS influences its interaction with TRF2 and dictates the repair of telomeres (57). RAD50 downregulates the association of TRF1 from telomeres and also contribute to maintain telomere length (58). We saw a decrease in telomerase activity across miRs and siRNAs experiments targeting the MRN complex. Moreover, the MRE11a target protector largely restored this function, implying this was the miR-494 target responsible for the compromised telomerase activity. Therefore, we suspect that the telomere maintenance function of the MRN complex drives the EC senescence downstream of miR-494 and miR-99b induced by DNA damage.

Our observations indicate that the function of these miRs in regulating telomerase activity and EC senescence may depend on the expression of VEGFR2. This is surprising, since expression of VEGFR2 may steer cells into a robust replication program rather than a cell cycle exit characteristic of senescence. However, it is possible that replication stress and DNA damage are enhanced in the presence of these miRs. Similarly, our findings demonstrate a decrease in MRN activity either by miR-99b induction or Mirin-1 affects the VEGFR signaling at the level of pERK. These data argue that the MRN pathway and the VEGFR2 signaling cross-talk functions as a threshold to determine the level of DNA damage in ECs. In the presence of significant DNA damage the MRN pathway is disrupted and the VEGF signaling is diminished thereby decreasing angiogenesis.

It is possible that during physiological aging, increase in miR-494 and miR-99b levels render the ECs less responsive to VEGF levels and therefore, decrease angiogenic responses. We envision that this senescence associated decrease in VEGF sensitivity functions in a feedforward loop and contributes to cardiovascular aging. Data from our mouse models demonstrates the utility of exploiting this pathway to decrease pathological angiogenesis. On the basis of these studies, we propose that the MRN complex is novel anti-angiogenic target.

## Materials and Methods

### Cell Culture and Reagents

HUVECs and HMVECs (Lonza) were cultured in EGM-2 media (Lonza) supplemented with 10% Fetal Calf Serum (Hyclone). NHLFs (Lonza) were cultured in FBM media (Lonza) supplemented with 10% Fetal Calf Serum (Hyclone). 4T1 cells (ATCC) were culture in McCoy’s media, DMEM or RPMI-1640 supplemented with 10% Fetal Calf Serum and antibiotics. Cells were tested and found negative for mycoplasma contamination before use in the assays described. Mirin-1 and Vandetanib were purchased from Cayman Chemical and Selleckchem respectively. VEGF was purchased from PeproTech, Inc.

### miRs/anti-miRs/siRNAs

miR mimics, inhibitors and respective controls were purchased from Life Technologies and Exiqon. For *in vivo* studies, high-performance liquid chromatography-purified miRs were purchased from Life Technologies in bulk quantities. siRNAs against MRE11a, RAD50 and NBN were purchased from Life Technologies. VEGFR2 Gapmer and MRE11a target protector were purchased from Exiqon.

### Vectors/Plasmids

MRE11a Luciferase-3-UTR plasmid was purchased from SwitchGear Genomics. RAD50 and NBN luciferase constructs were generated by cloning the entire 3’UTR regions into pmiR-REPORT vector (Ambion). Luciferase assay reagents were purchased from SwitchGear Genomics and Promega.

### Transfection

Cell were transfected at 50-60% confluence using standard forward transfection protocols using RNAimax reagent (Life Technologies) for miRs, siRNAs or gapmers and Lipofectamine 2000 for plasmid or plasmid RNA dual transfections. Typically 50 nM RNA and 1-2 μ g plasmid DNA were used for transfections. Target protectors were transfected at a concentration of 50 nM or equivalent to the miR amounts.

### Radiation of Cells/Mice

Cells or mice were irradiation on a Shepherd□137cesium irradiator at a rate of ~166 cGy/min. In tumor-targeted radiation experiments, mice were restrained in a lead shield with flank cut-outs (Brain tree scientific) to minimize exposure to the non-tumor areas.

### β-Gal Senescence assay

HUVEC were transfected for 48-72h with the microRNAs or siRNAs. After this time cells were washed with cold PBS and then stained for β-galactosidase activity following manufacturer’s protocol (Senescence Cells Histochemical Staining Kit, Sigma).

### Telomerase activity assay

Cells were transfected with microRNAs or siRNAs for 24h. Cells were lysed and processed according to manufacturer’s instructions (Quantitative Telomerase Detection Kit, Allied Biotech). The telomerase activity level in the cell extract was determined through its ability to synthesize telomeric repeats onto an oligonucleotide substrate. The resultant extended product was subsequently amplified by polymerase chain reaction (PCR).

### Cell Titer Glo/ Caspase Glo

Cells were transfected in a 6 well plate with miR mimics, siRNAs or inhibitors, and the corresponding controls from Life Technologies as previously described (59)‥ Cell Titer-Glo and Caspase 3/7 Glo were analyzed at 48-96h post treatments, according to manufacturer’s instructions.

### Western blot and densitometric analysis

After treatment, cells were washed in phosphate-buffered saline (PBS) and lysed in RIPA buffer (Sigma) supplemented with Complete Protease inhibitor cocktail (ROCHE) and Phosphatase inhibitors cocktail 2 and 3 (Sigma). Lysed cells were harvested by scraping, and proteins were analyzed by Western blot. Equivalent amounts of protein were loaded on a 4-12% gradient SDS-polyacrylamide gel (BioRAD) and transferred for 30 min in a TransBlot turbo (BioRAD) onto Nitrocellulose membranes. Membranes were blocked in 5% milk or 3% BSA and incubated with antibodies as indicated: MRE11a (Cell Signaling, 4847, 1:1000), RAD50 (Cell Signaling, 3427, 1:1000), NBS1 (Cell Signaling, 14956, 1:500), p21 (Cell Signaling, 2947, 1:1000), pRb (Cell Signaling, 9301, 1:500),. β-actin (Sigma, A5316, 1:10,000 1□h RT) was used as a housekeeping control for the total levels of protein loaded. Membranes were washed in TBST and incubated with secondary antibodies from Licor Biosciences. Licor antibodies used were goat anti mouse 925-68020 (1:15,000) and goat anti rabbit 925-32211 (1:15,000). Blots were scanned on the Licor Odyssey scanner according to manufacturer’s instructions.

### RNA extraction, RT-PCR, miR Profiling

Total RNA and microRNA were isolated using a miRVana microRNA isolation kit (Ambion). Reverse transcription was performed using TaqMan™ Advanced cDNA Synthesis Kit (Life Tech) according to the manufacturer’s instructions. RT-PCR was performed using multiplexed TaqMan primers (Applied Biosystems). The relative quantification of gene expression was determined using the 2^-△△Ct^ method (35). Using this method, we obtained the fold changes in gene expression normalized to an internal control gene, GAPDH or U6 snRNA, respectively. For target analysis, a 92 gene DNA damage array (Life tech, 4418773) was used and for senescence phenotype profiling, the Human Cellular Senescence array was utilized (SA Biosciences) per manufacturer’s recommendations.

### vH2AX staining

100,000 HUVECs were cultured on glass coverslips in 24-well plates and transfected with miRs/siRNAs using RNAimax (Life Technologies). Cells were fixed at different time points with 4% paraformaldehyde for 10 minutes at room temperature, permeabilized with 90% methanol for 10 minutes at 4°C.□□Coverslips were blocked with 1.5% normal goat serum (NGS) and incubated with primary antibody H2AX (Abcam, 11174, 1:500), at a 1:1000 dilution in NGS for 1h, washed and then incubated with secondary antibody for 30 minutes, washed and then mounted on glass slides for confocal imaging.

### miR in-situ hybridization

In-situ hybridization was performed on frozen tumor sections as described in our previous studies (60, 61) using a DIG labeled miR-494 Locked Nucleic Acid (LNA) probe (Exiqon). DIG was detected by an anti-DIG HRP antibody (Roche) and amplified using a TSA-Plus Cy3 system (Perkin Elmer).

### miR-TRAP/RISC TRAP assay

293 T cells were co-transfected with a plasmid coding for a flag-tagged dominant negative GW418 mutant (Clontech kit #632016) along with a control mimic, miR-99b or miR-494 mimic according to kit instructions. 24h later the RNA protein complexes were crosslinked and the RISC complex was immunoprecipitated using an anti-FLAG antibody and RNA was isolated for quantitative real-time PCR of target genes. The fold enrichment was calculated using pre and post IP controls as well as normalization to the control mimic pull-downs.

### 3-D Angiogenic Sprouting Assay

Early passage HUVECs were coated on cytodex-3 beads (GE Healthcare) at a density of 10 million cells/40 μl beads and incubated in suspension for 3-4 hours with gentle mixing every hour. They were plated on TC treated 6 well dishes overnight and resuspended in a 2mg/ml fibrin gel with 200,000 human smooth muscle cells. The gel was allowed to polymerize and complete EGM-2 media was added. Sprouts were visualized from days 3-4 via confocal imaging after overnight incubation with FITC labeled *Ulex europaeus* lectin (Vector labs). Immunofluorescence imaging was performed on a Yokogawa CSU-W1 spinning disk confocal microscope with 20x, 0.45 Plan Fluor objective (Nikon).

### Flow cytometry

CD44 expression was analyzed by flow cytometry in HUVECs transfected with miR-494 for 48 hours. Cells were washed in PBS and trypsinized. Cells were incubated in blocking solution (0.5 %BSA/ 10% goat serum) for 30 min and then incubated for 1 hour in primary antibody-PEcy7 conjugated (CD44, BD, 560533) prepared in blocking solution. Then cells were washed 3X with PBS. Cells were analyzed in a CANTO II flow cytometer.

### Cell cycle analysis

HUVEC were transfected for 48 hours with microRNAs, inhibitors or siRNAs. Cells were then harvested, washed, and fixed in 70% ice-cold ethanol at 4°C overnight. Then, cells were centrifuged, washed with cold PBS, and re-centrifuged. Cells were then re-suspended in 250□μL PBS and stained with 10□μL propidium iodide (PI, 1 Cmg/mL) and 10 μL RNase A (10□mg/mL) for 30□min at room temperature. DNA content was assessed using flow cytometry (CANTO II) to calculate the percentage of cells in subG1, G0/G1, S, and G2/M phases with FlowJo software.

### Immunofluorescence and microscopy

In some experiments, CD31a and MRE11a were visualized using immunofluorescence staining from OCT sections. Slides were fixed with 4% PFA and stained overnight for CD31 488 (BD bioscience 611986 1:200 o/n), MRE11a (Cell Signaling Technology 1:100) and Phalloidin Alexa 647 (1:50). For MRE11a antibody, following day fixed cells were incubated with Goat anti-Rabbit Alexa 488 (1:500) in 5%BSA/TBS for 1 hour. Imaging was performed on a Nikon Spectral C1 confocal microscope (Nikon C1si with EZC1 acquisition software, Nikon Instruments) with Plan Apo 10X/0.45 air, Plan Apo 20X/0.75 air, and Plan Apo 60X/1.40 oil objective lenses (Nikon). Some immunofluorescence imaging was performed on a Yokogawa CSU-W1 spinning disk confocal microscope with 20 0.45 Plan Fluor objective (Nikon). All images were taken with a channel series. Images were analyzed with Image J software for quantitation.

### pERK ELISA

We measured the abundance of p-ERK with an ELISA kit (PathScan^®^ Phospho-p44/42 MAPK (Thr202/Tyr204) Sandwich ELISA, Cell Signaling Technology) following manufacturer’s instructions.

### *In vivo* assays

All animal work was approved by the OHSU Institutional Animal Use and Care Committee. Immune-compromised 8-10 week old female nu/nu mice were purchased from Jackson Labs. Growth factor reduced Matrigel (BD) with 400 ng/ml recombinant human bFGF (Millipore) was injected subcutaneously in nu/nu mice. Mice were injected i.v. with 7C1-nanoparticles containing miR-494 or control miR (~1mg/kg, i.v) 3 or 4 days after plugs were implanted. At day 7 mouse tissues were harvested and processed to obtain RNA or frozen in OCT for tissue staining. 4T1 cells (1X10s^4) were implanted into the mammary fat pad #4 of 6-8 week old female Balb/C mice in 100μl Matrigel. Mice were randomized into groups once the average tumor volume reached 150 mm^3^, approximately 10 days after implantation. Mice were treated with 7C1-nanoparticles containing miR-103 or control miR (0.7 mg/kg, i.v.). Mice were euthanized ~ day 18-20 for analysis of metastatic burden in lungs. Additional experiments were carried out in postpubertal female FVB/n mice (10-12 weeks), which received injections of 50,000 PyMT-derived tumor cells into the right lower mammary fat pad for induction of orthotopic tumors as described in (62).

### Statistics

All statistical analysis was performed using Excel (Microsoft) or Prism (GraphPad). Two-tailed Student’s t-test was used to calculate statistical significance. Variance was similar between treatment groups.

## Acknowledgements

We thank Lisa Coussens (OHSU) for help with the PyMT model and sharing archived FFPE tissue sections from (62). We thank Sergio Fazio and Stephen Lloyd (OHSU) for useful discussions. We thank Shushan Rana and Katherine Kelly (OHSU) for comments on the manuscript. We acknowledge the OHSU Advanced Light Microscopy Core, Knight Cancer Institute Flow Cytometry Core and the Gene Profiling Shared Resource for technical help and useful discussions. This work was supported by US NIH grant R00HL112962, R56HL137779 and an innovative research grant from the American Heart Association (17IRG33400218) all to S.A.

---

**Supplementary Figure 1.**
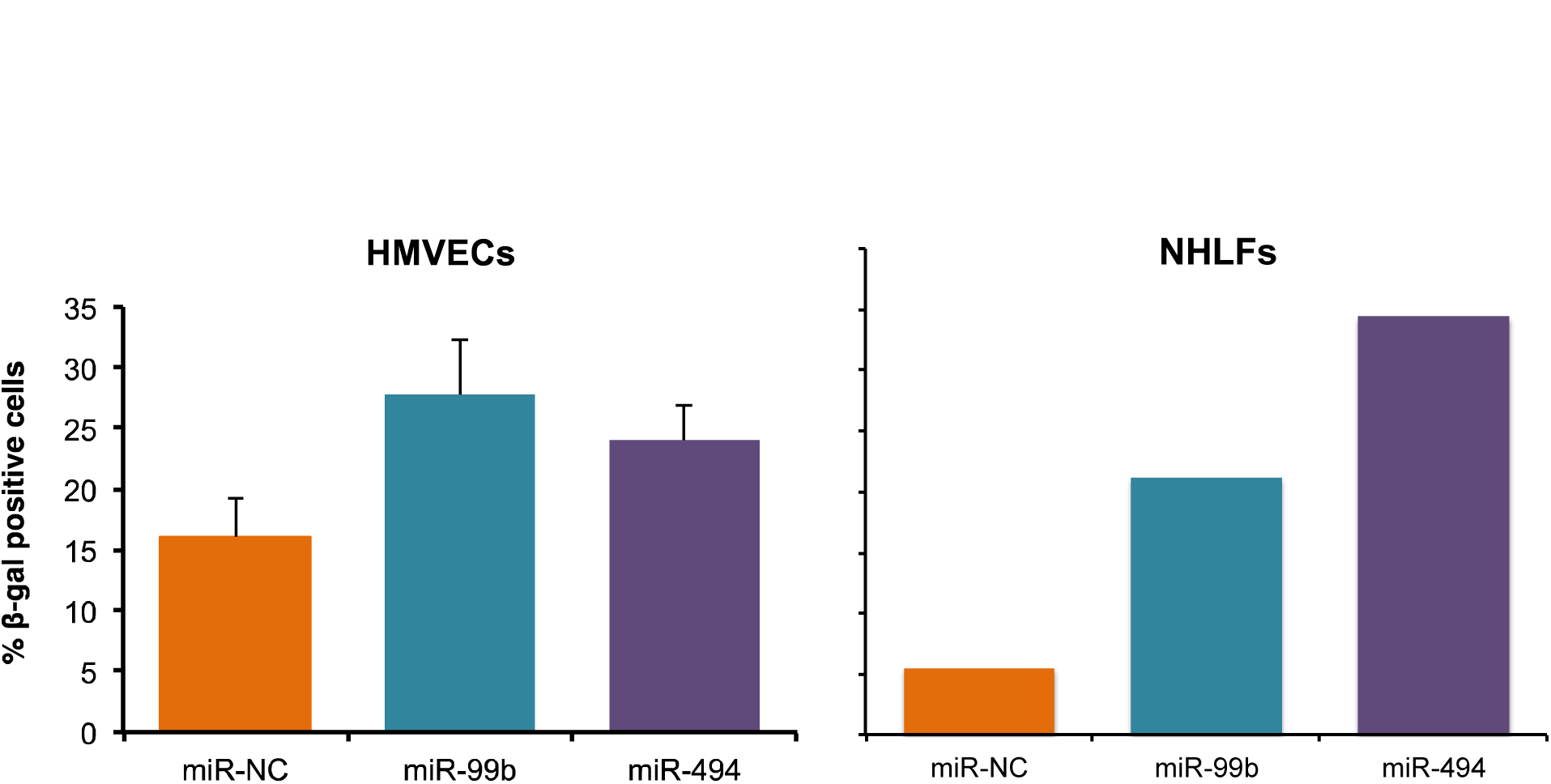
Senescence phenotype in other normal cells. β-Gal assay in HMVECs or NHLFs transfected for 48 hours with miR-C, miR-99b or miR-494. Bars show % mean ± SEM of β-gal positive cells for at least hundred cells analyzed.

**Supplementary Figure 2.**
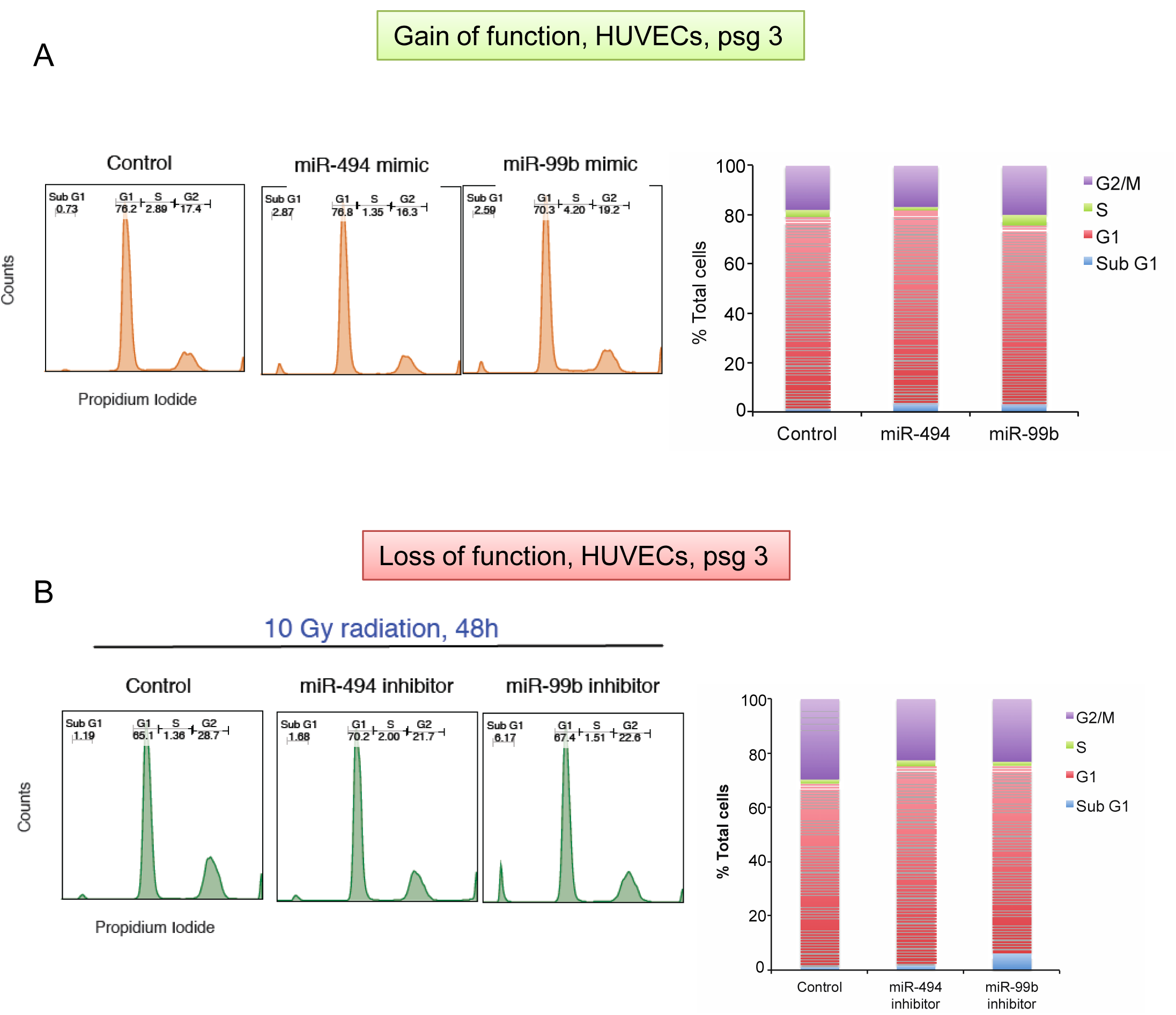
Gain and loss of senescence miRs affects cell cycle progression. HUVECs were transfected with the indicated miR mimics (A) or inhibitors (B) and cell cycle analysis was performed via flow cytometry of propidium iodide stained cells at 48h post transfection (A). Transfected cells were irradiated at 24h and cell cycle analysis was performed at 48h post radiation (B).

**Supplementary Figure 3.**
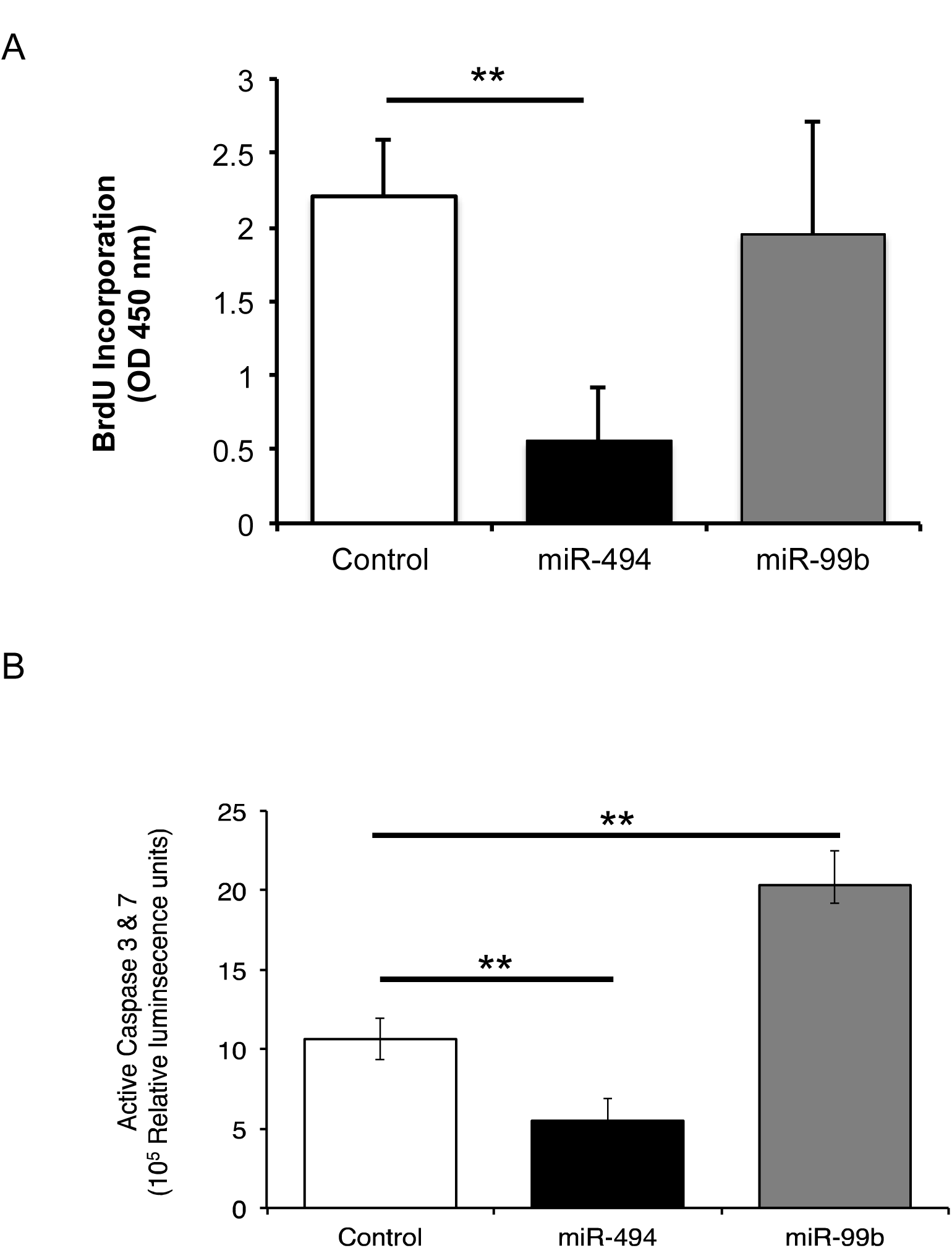
Gain and loss of senescence miRs affects proliferation and survival. HUVECs were transfected with the indicated miR mimics for 48h. Proliferation was measured by a BrdU incorporation assay following an overnight pulse of BrdU (A) and cell death was measured by a lumiscence based Caspase activity assay (B).

**Supplementary Figure 4.**
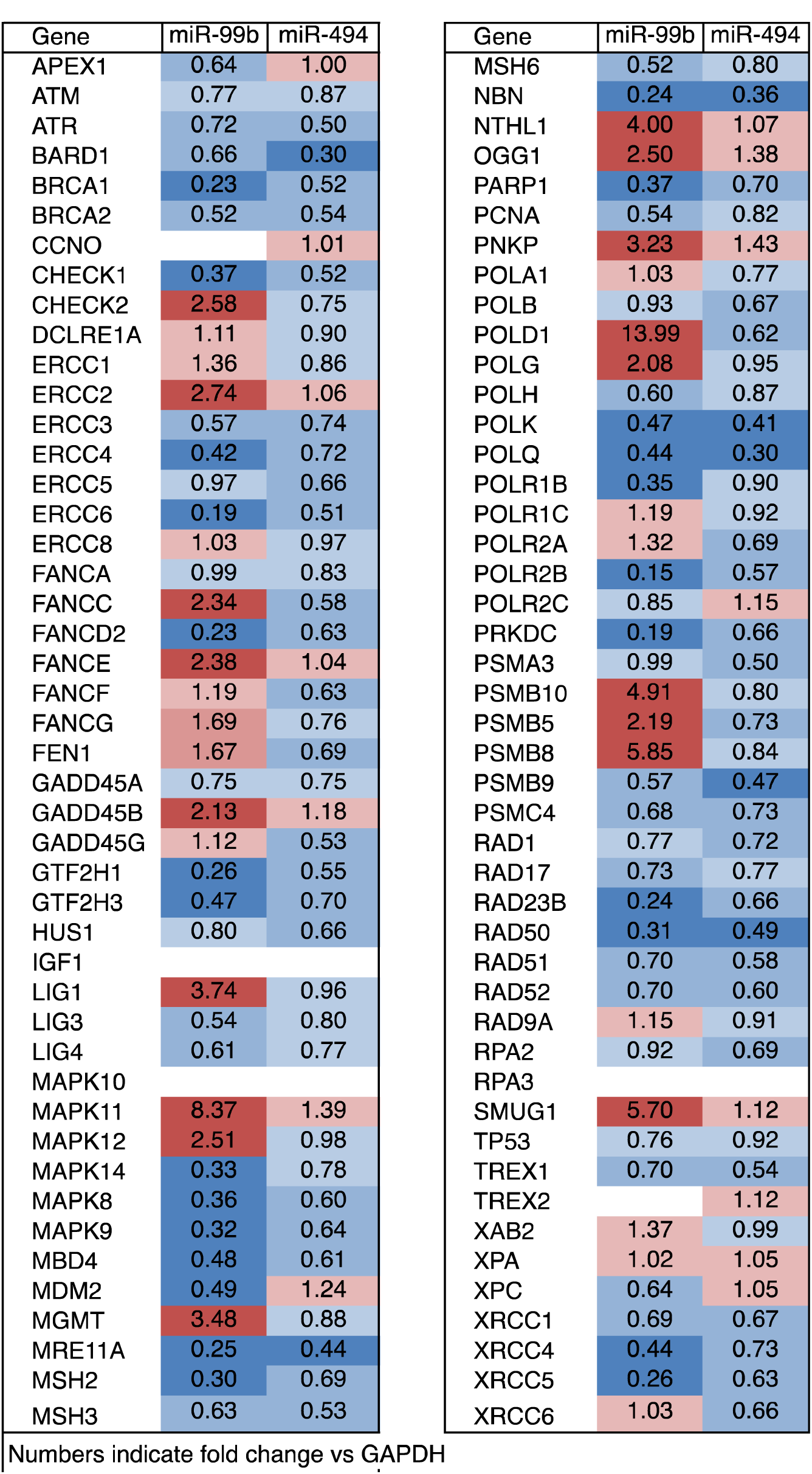
Identification of miR targets by targeted transcriptional profiling. Heat map depicting mRNA changes induced by miR-99b and miR-494 in HUVECs at 24h post transfection as analyzed by a TaqMan Human DNA Repair qRT-PCR gene signature array.

**Supplementary Figure 5.**
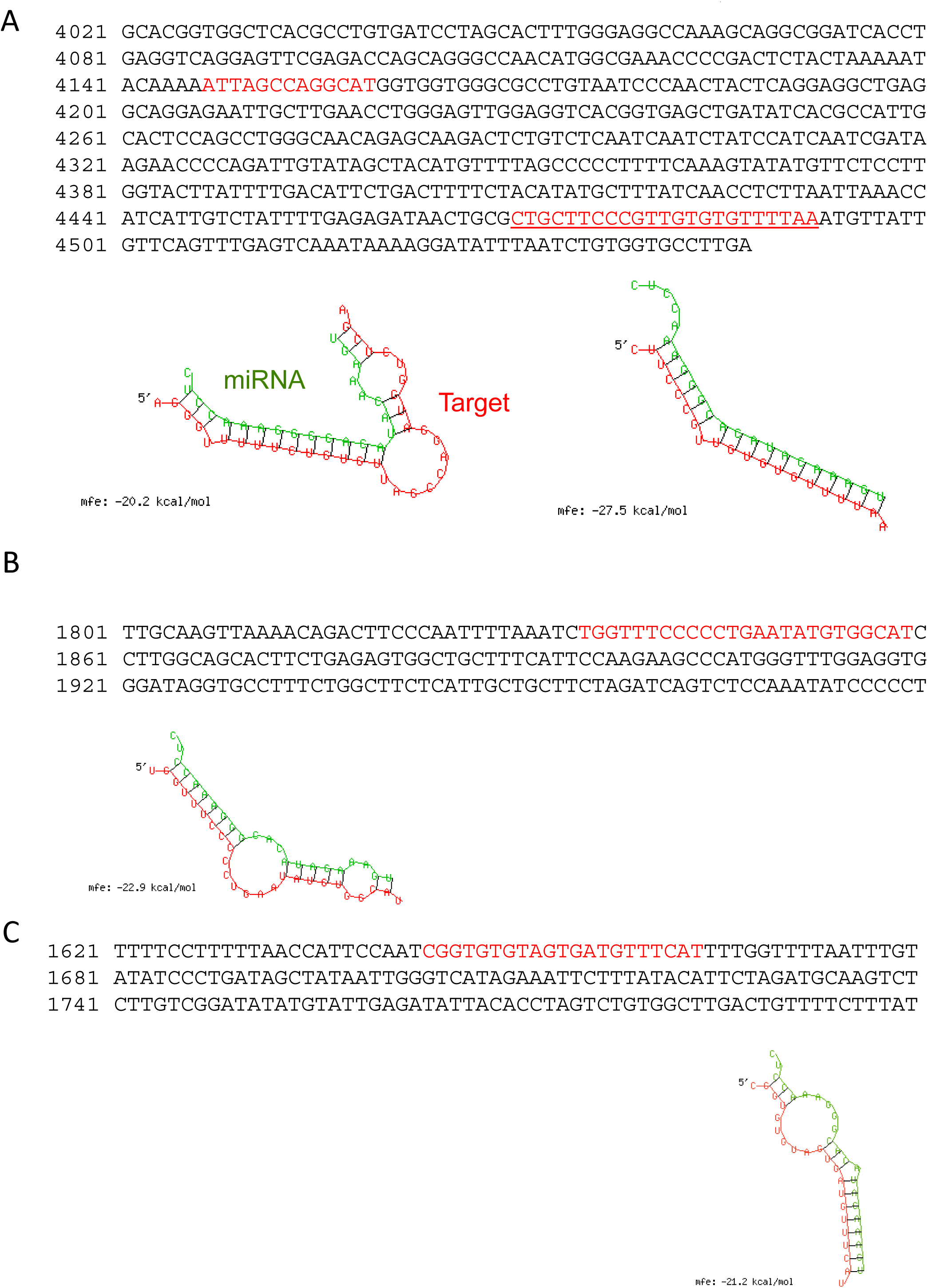
RNA hybrid models for miR-494. **A**. mRNA sequence of human MRE11a, **B**. RAD50 and **C**. NBN. Binding sites are highlighted in red. MRE11a-miR-494 target blocker was designed against the site highlighted in red and underlined.

**Supplementary Figure 6.**
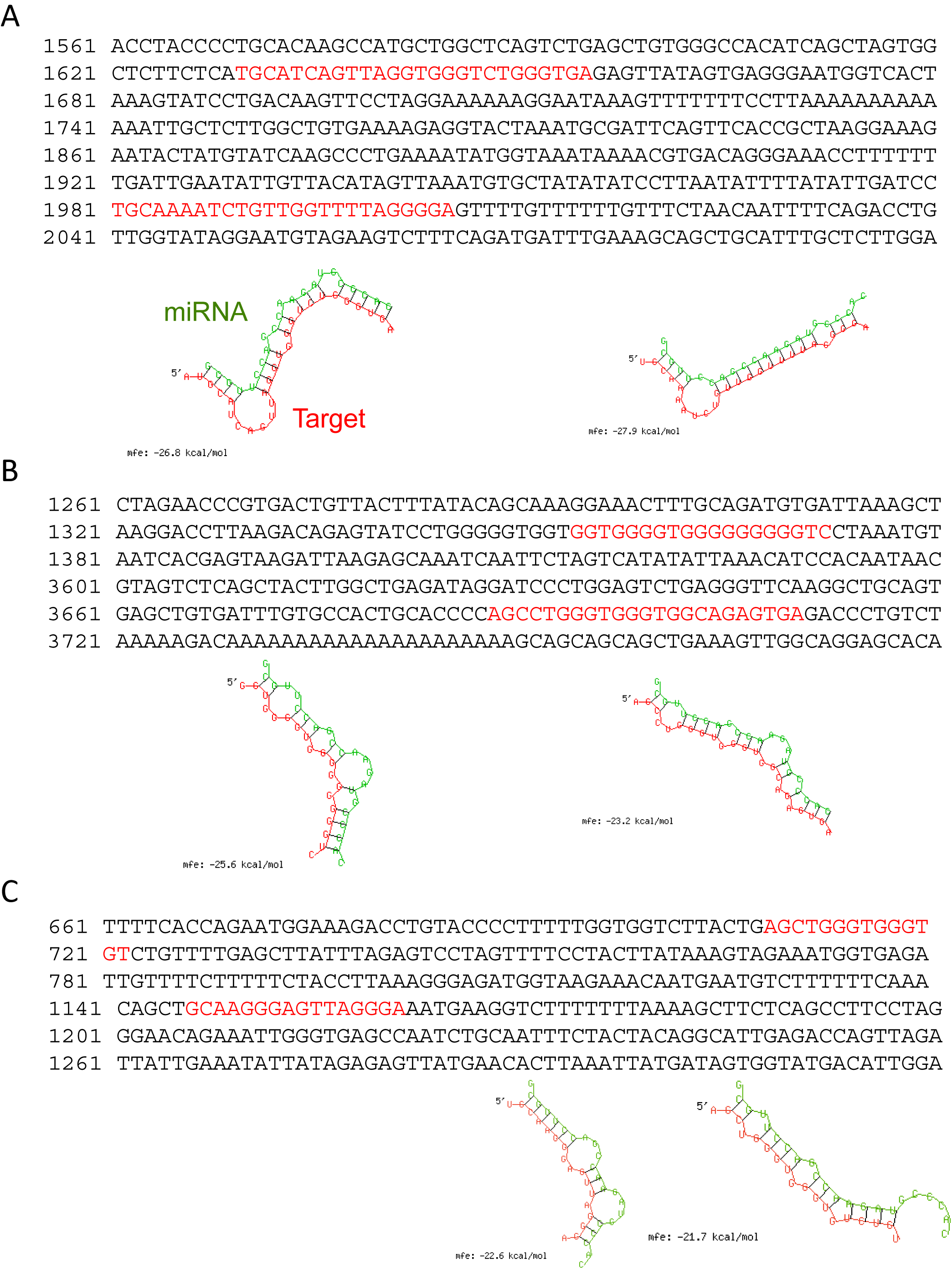
RNA hybrid models for miR-99b. **A**. mRNAsequence of human MRE11a **B**. RAD50 and **C**. NBN. Binding sites are highlighted in red.

**Supplementary Figure 7.**
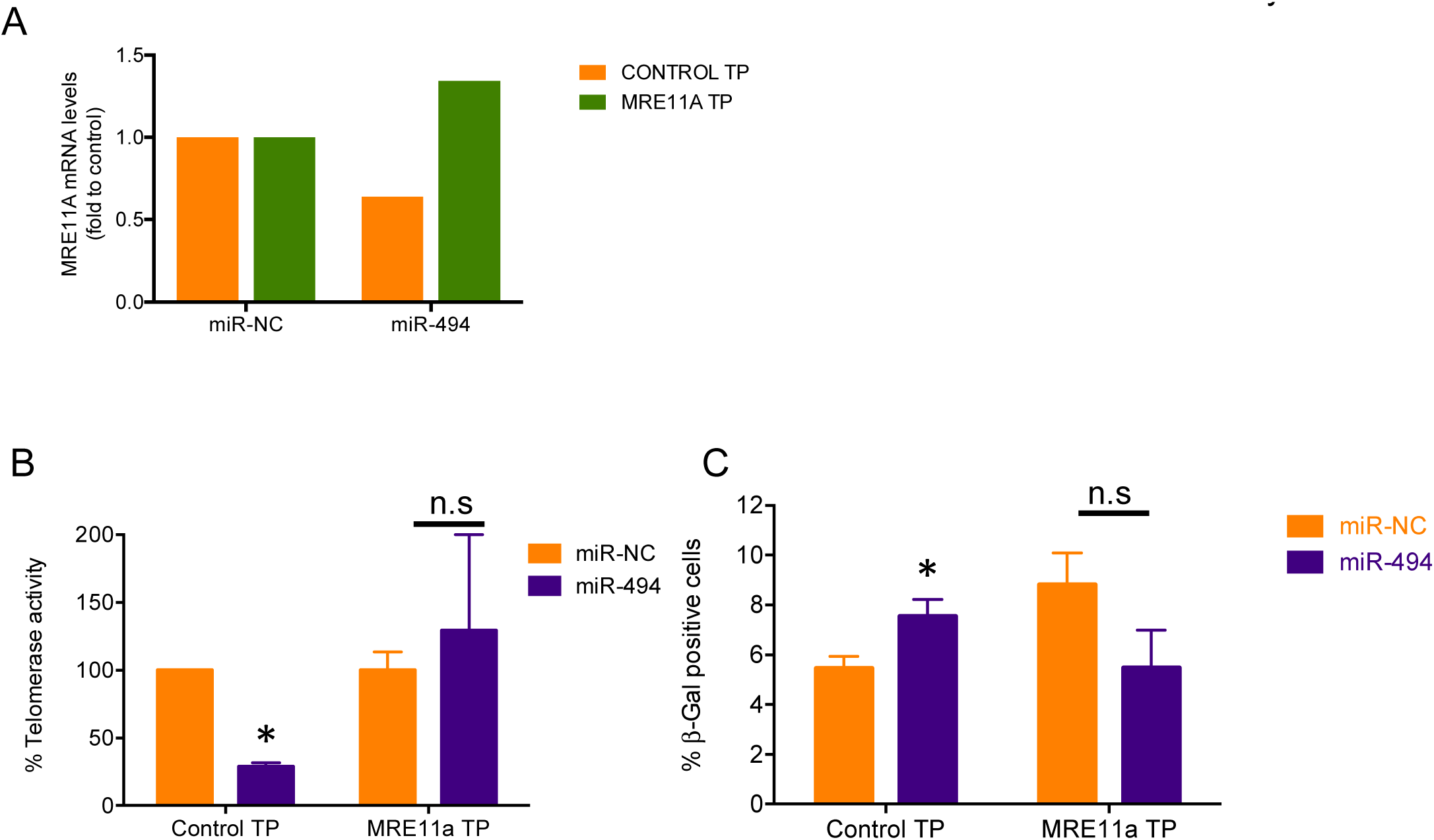
Protecting Mre11a from miR-494 rescues telomerase activity and decreases senescence. **A**. HUVECs were co-transfected with a target protector for miR-494 binding site in MRE11a-3’-UTR or a scrambled control target protector, and the corresponding miR. Bar graph depicts mean +SEM of Mre11a. **B-C**. HUVECs were transfected as described in A) for 48h and telomerase activity (B) and senescence associated β-Gal (C) was assayed as described in Fig 2. Bars depict % mean + SEM of Telomerase activity in B) and show % mean + SEM of β-gal positive cells for at least hundred cells analyzed in C).

**Supplementary Figure 8.**
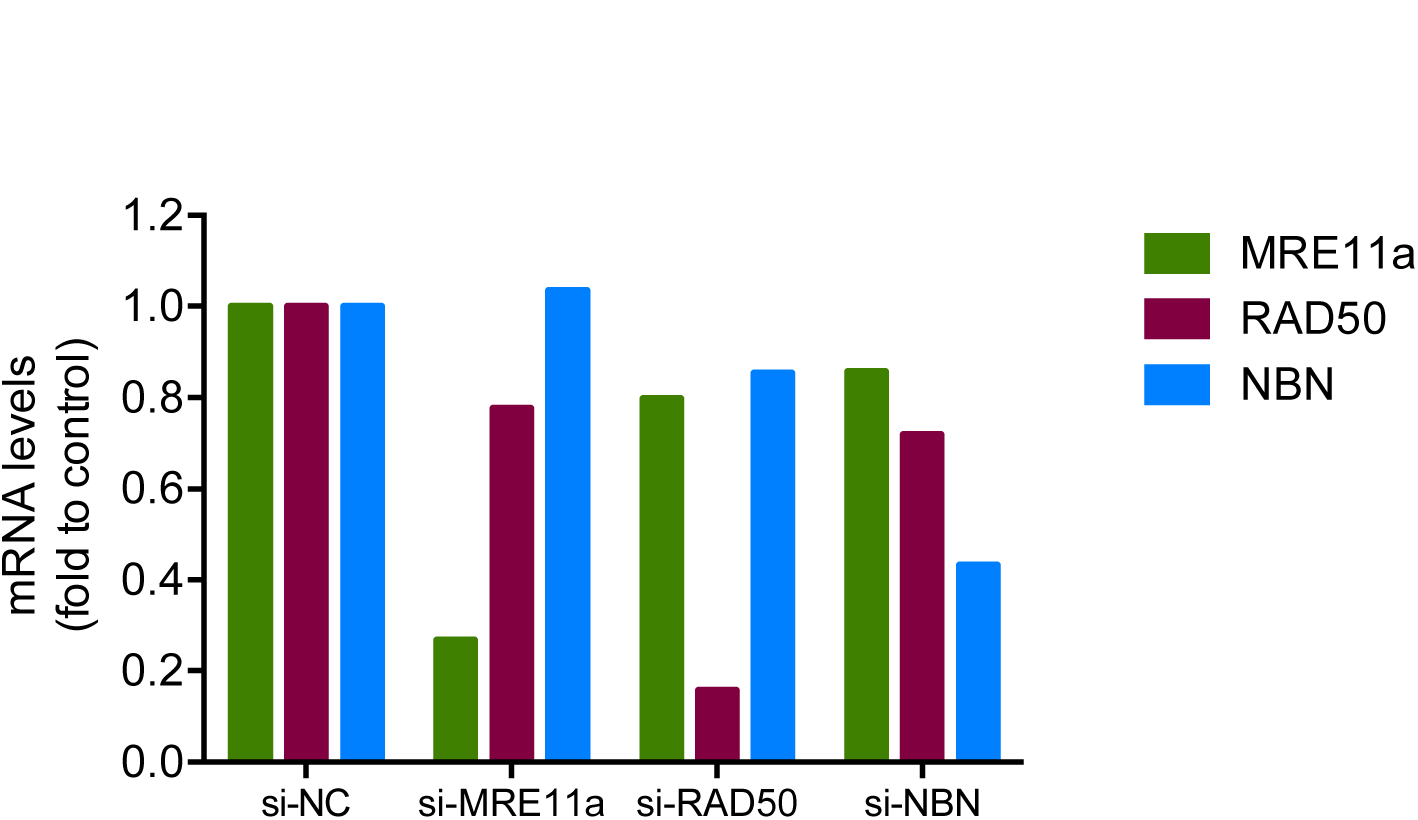
siRNA validation. HUVECs were transfected with either a negative control siRNA or specific siRNAs against MRE11a, RAD50 and NBN and qRT-PCR was performed at 24h for the indicated target genes.

**Supplementary Figure 9.**
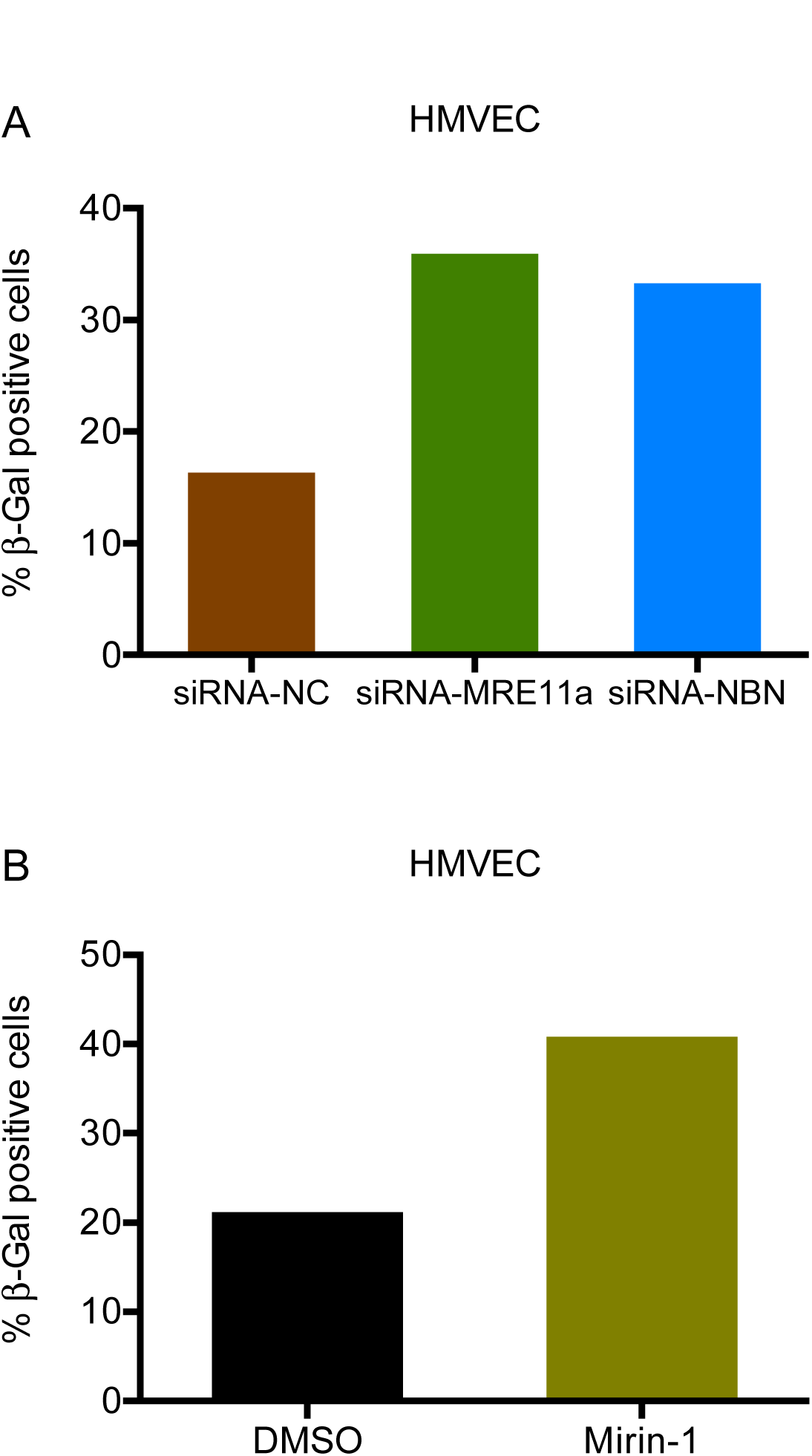
Senescence phenotype upon MRN disruption in HMVEC cells. β-Gal assay in HMVECs transfected for 48 hours with **A**. siRNA-NC, siRNA-MRE11a or siRNA-NBN or **B**. treated with Mirin-1 (50 μM) for 24h. Graphs show % mean ± SEM of β-gal positive cells for at least hundred cells analyzed.

**Supplementary Figure 10.**
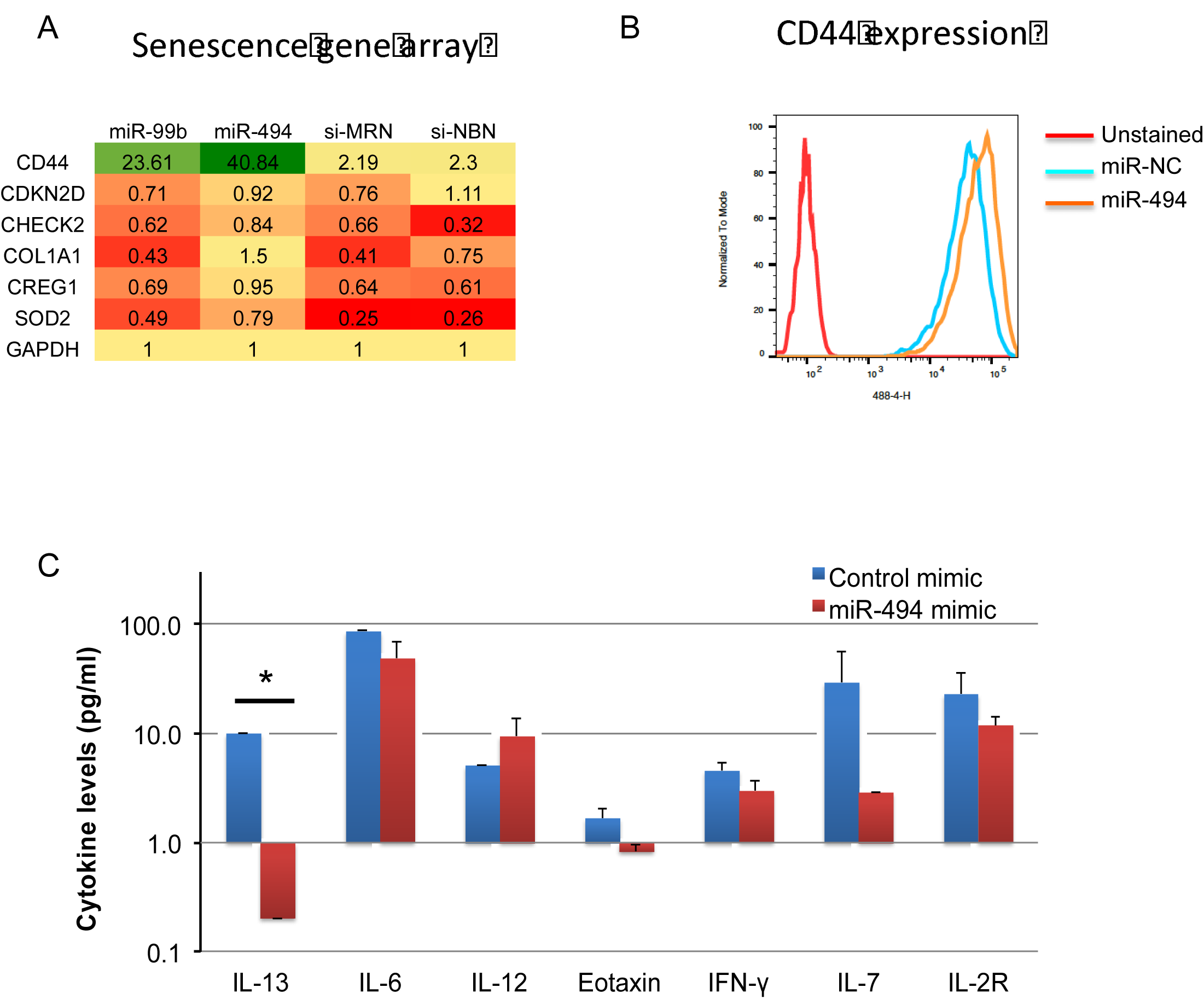
miR-494 regulates cell intrinsic and secreted proteins involved in senescence. **A**. Heat map represents the 6 genes that are similarly regulated by miR-99b, miR-494, siRNA-MRE11a and siRNA-RAD50 as analyzed by a qRT-PCR based Human Senescence gene signature Array. **B**. Representative CD44 histogram analyzed by Flow cytometry. HUVECs were transfected for 48 hours with miR-494 or a control miR. **C**. Supernatants from transfected cells at 48h was analyzed for differential expression of 24 cytokines. Cytokines that were altered at least 1.5 fold are depicted. Bars depict mean + SEM. * P<0.001 by Student’s T-test.

## References

1. Bautista-Nino PK, Portilla-Fernandez E, Vaughan DE, Danser AH, Roks AJ. DNA Damage: A Main Determinant of Vascular Aging. Int J Mol Sci. 2016 May 18;17(5). PubMed PMID: 27213333. Pubmed Central PMCID: PMC4881569.

2. Regina C, Panatta E, Candi E, Melino G, Amelio I, Balistreri CR, et al. Vascular ageing and endothelial cell senescence: Molecular mechanisms of physiology and diseases. Mech Ageing Dev. 2016 Oct;159:14–21. PubMed PMID: 27155208.

3. Gates PE, Strain WD, Shore AC. Human endothelial function and microvascular ageing. Exp Physiol. 2009 Mar;94(3):311–6. PubMed PMID: 19042980.

4. Economopoulou M, Langer HF, Celeste A, Orlova VV, Choi EY, Ma M, et al. Histone H2AX is integral to hypoxia-driven neovascularization. Nat Med. 2009 05//print;15(5):553–8.

5. Okuno Y, Nakamura-Ishizu A, Otsu K, Suda T, Kubota Y. Pathological neoangiogenesis depends on oxidative stress regulation by ATM. Nat Med. 2012 08//print;18(8):1208–16.

6. Guo Z, Kozlov S, Lavin MF, Person MD, Paull TT. ATM Activation by Oxidative Stress. Science. 2010;330(6003):517–21.

7. Kang HT, Park JT, Choi K, Kim Y, Choi HJC, Jung CW, et al. Chemical screening identifies ATM as a target for alleviating senescence. Nat Chem Biol. 2017 06//print;13(6):616–23.

8. Williams RS, Williams JS, Tainer JA. Mre11-Rad50-Nbs1 is a keystone complex connecting DNA repair machinery, double-strand break signaling, and the chromatin template. Biochem Cell Biol. 2007 Aug;85(4):509–20. PubMed PMID: 17713585.

9. Lafrance-Vanasse J, Williams GJ, Tainer JA. Envisioning the dynamics and flexibility of Mre11-Rad50-Nbs1 complex to decipher its roles in DNA replication and repair. Prog Biophys Mol Biol. 2015 Mar;117(2-3):182–93. PubMed PMID: 25576492. Pubmed Central PMCID: PMC4417436.

10. Dimitrova N, de Lange T. Cell cycle-dependent role of MRN at dysfunctional telomeres: ATM signaling-dependent induction of nonhomologous end joining (NHEJ) in G1 and resection-mediated inhibition of NHEJ in G2. Mol Cell Biol. 2009 Oct;29(20):5552–63. PubMed PMID: 19667071. Pubmed Central PMCID: PMC2756883.

11. Porro A, Feuerhahn S, Lingner J. TERRA-reinforced association of LSD1 with MRE11 promotes processing of uncapped telomeres. Cell Rep. 2014 Feb 27;6(4):765–76. PubMed PMID: 24529708.

12. Ju YJ, Lee KH, Park JE, Yi YS, Yun MY, Ham YH, et al. Decreased expression of DNA repair proteins Ku70 and Mre11 is associated with aging and may contribute to the cellular senescence. Exp Mol Med. 2006 Dec 31;38(6):686–93. PubMed PMID: 17202845.

13. Gao R, Singh R, Kaul Z, Kaul SC, Wadhwa R. Targeting of DNA Damage Signaling Pathway Induced Senescence and Reduced Migration of Cancer cells. J Gerontol A Biol Sci Med Sci. 2015 Jun;70(6):701–13. PubMed PMID: 24747666.

14. Boon RA, Dimmeler S. MicroRNAs in myocardial infarction. Nat Rev Cardiol. 2015 Mar;12(3):135–42. PubMed PMID: 25511085.

15. Santulli G. MicroRNAs and Endothelial (Dys) Function. J Cell Physiol. 2016 Aug;231(8):1638–44. PubMed PMID: 26627535. Pubmed Central PMCID: PMC4871250.

16. Fish JE, Srivastava D. MicroRNAs: opening a new vein in angiogenesis research. Science signaling. 2009 Jan 6;2(52):pe1. PubMed PMID: 19126861. Pubmed Central PMCID: 2680274.

17. Shi H, Li P, Liang W, Chen J, Gao Y. Mechanisms of microRNA-mediated regulation of angiogenesis. Frontiers in bioscience. 2010;2:1304–19. PubMed PMID: 20515803.

18. Landskroner-Eiger S, Moneke I, Sessa WC. miRNAs as modulators of angiogenesis. Cold Spring Harbor perspectives in medicine. 2013 Feb;3(2):a006643. PubMed PMID: 23169571. Pubmed Central PMCID: 3552340.

19. Bartel DP. MicroRNAs: target recognition and regulatory functions. Cell. 2009 Jan 23;136(2):215–33. PubMed PMID: 19167326. Pubmed Central PMCID: PMC3794896.

20. Ebert MS, Sharp PA. Roles for microRNAs in conferring robustness to biological processes. Cell. 2012;149(3):515–24. PubMed PMID: PMC3351105.

21. Osella M, Bosia C, Corá D, Caselle M. The Role of Incoherent MicroRNA-Mediated Feedforward Loops in Noise Buffering. PLoS Computational Biology. 2011 03/10 04/03/received 01/28/accepted;7(3):e1001101. PubMed PMID: PMC3053320.

22. Li X, Cassidy JJ, Reinke CA, Fischboeck S, Carthew RW. A MicroRNA Imparts Robustness against Environmental Fluctuation during Development. Cell. 2009 4/17/;137(2):273–82.

23. Inui M, Martello G, Piccolo S. MicroRNA control of signal transduction. Nat Rev Mol Cell Biol. 2010 04//print;11(4):252–63.

24. Garner KM, Pletnev AA, Eastman A. Corrected structure of mirin, a small-molecule inhibitor of the Mre11-Rad50-Nbs1 complex. Nat Chem Biol. 2009 Mar;5(3):129–30; author reply 30. PubMed PMID: 19219009. Pubmed Central PMCID: PMC3881006.

25. Ponta H, Sherman L, Herrlich PA. CD44: From adhesion molecules to signalling regulators. Nat Rev Mol Cell Biol. 2003 01//print;4(1):33–45.

26. Tremmel M, Matzke A, Albrecht I, Laib AM, Olaku V, Ballmer-Hofer K, et al. A CD44v6 peptide reveals a role of CD44 in VEGFR-2 signaling and angiogenesis. Blood. 2009;114(25):5236–44.

27. Corne J, Chupp G, Lee CG, Homer RJ, Zhu Z, Chen Q, et al. IL-13 stimulates vascular endothelial cell growth factor and protects against hyperoxic acute lung injury. The Journal of Clinical Investigation. 2000 09/15/;106(6):783–91.

28. Kim HD, Yu S-J, Kim HS, Kim Y-J, Choe JM, Park YG, et al. Interleukin-4 Induces Senescence in Human Renal Carcinoma Cell Lines through STAT6 and p38 MAPK. The Journal of biological chemistry. 2013 08/09 07/11/received;288(40):28743–54. PubMed PMID: PMC3789971.

29. Wilson R, Espinosa-Diez C, Kanner N, Chatterjee N, Ruhl R, Hipfinger C, et al. MicroRNA regulation of endothelial TREX1 reprograms the tumour microenvironment. Nat Commun. 2016 Nov 25;7:13597. PubMed PMID: 27886180. Pubmed Central PMCID: 5133658 modulate blood vessel growth, patterning, tumour growth and malignant disease and method for making and using them. The remaining authors declare no competing financial interests.

30. Dahlman JE, Kauffman KJ, Langer R, Anderson DG. Nanotechnology for in vivo targeted siRNA delivery. Adv Genet. 2014;88:37–69. PubMed PMID: 25409603.

31. Zhao JJ, Yang J, Lin J, Yao N, Zhu Y, Zheng J, et al. Identification of miRNAs associated with tumorigenesis of retinoblastoma by miRNA microarray analysis. Child’s nervous system: ChNS: official journal of the International Society for Pediatric Neurosurgery. 2009 Jan;25(1):13–20. PubMed PMID: 18818933.

32. Wang X, Zhang X, Ren XP, Chen J, Liu H, Yang J, et al. MicroRNA-494 targeting both proapoptotic and antiapoptotic proteins protects against ischemia/reperfusion-induced cardiac injury. Circulation. 2010 Sep 28;122(13):1308–18. PubMed PMID: 20837890. Pubmed Central PMCID: 2956502.

33. Wezel A, Welten SM, Razawy W, Lagraauw HM, de Vries MR, Goossens EA, et al. Inhibition of MicroRNA-494 Reduces Carotid Artery Atherosclerotic Lesion Development and Increases Plaque Stability. Annals of surgery. 2015 Nov;262(5):841–8. PubMed PMID: 26583674.

34. Asuthkar S, Velpula KK, Nalla AK, Gogineni VR, Gondi CS, Rao JS. Irradiation-induced angiogenesis is associated with an MMP-9-miR-494-syndecan-1 regulatory loop in medulloblastoma cells. Oncogene. 2014 Apr 10;33(15):1922–33. PubMed PMID: 23728345.

35. Chen S, Zhao G, Miao H, Tang R, Song Y, Hu Y, et al. MicroRNA-494 inhibits the growth and angiogenesis-regulating potential of mesenchymal stem cells. FEBS letters. 2015 Mar 12;589(6):710–7. PubMed PMID: 25660325.

36. Welten SM, Bastiaansen AJ, de Jong RC, de Vries MR, Peters EA, Boonstra MC, et al. Inhibition of 14q32 MicroRNAs miR-329, miR-487b, miR-494, and miR-495 increases neovascularization and blood flow recovery after ischemia. Circulation research. 2014 Sep 26;115(8):696–708. PubMed PMID: 25085941.

37. Esser JS, Saretzki E, Pankratz F, Engert B, Grundmann S, Bode C, et al. Bone morphogenetic protein 4 regulates microRNAs miR-494 and miR-126-5p in control of endothelial cell function in angiogenesis. Thromb Haemost. 2017 Jan 26. PubMed PMID: 28124060.

38. Comegna M, Succoio M, Napolitano M, Vitale M, D’Ambrosio C, Scaloni A, et al. Identification of miR-494 direct targets involved in senescence of human diploid fibroblasts. FASEB J. 2014 Aug;28(8):3720–33. PubMed PMID: 24823364.

39. Weng JH, Yu CC, Lee YC, Lin CW, Chang WW, Kuo YL. miR-494-3p Induces Cellular Senescence and Enhances Radiosensitivity in Human Oral Squamous Carcinoma Cells. Int J Mol Sci. 2016 Jul 08;17(7). PubMed PMID: 27399693. Pubmed Central PMCID: PMC4964468.

40. Ohdaira H, Sekiguchi M, Miyata K, Yoshida K. MicroRNA-494 suppresses cell proliferation and induces senescence in A549 lung cancer cells. Cell Prolif. 2012 Feb;45(1):32–8. PubMed PMID: 22151897.

41. Liu Y, Li X, Zhu S, Zhang JG, Yang M, Qin Q, et al. Ectopic expression of miR-494 inhibited the proliferation, invasion and chemoresistance of pancreatic cancer by regulating SIRT1 and c-Myc. Gene Ther. 2015 Sep;22(9):729–38. PubMed PMID: 25965392.

42. Kane NM, Howard L, Descamps B, Meloni M, McClure J, Lu R, et al. Role of microRNAs 99b, 181a, and 181b in the differentiation of human embryonic stem cells to vascular endothelial cells. Stem Cells. 2012 Apr;30(4):643–54. PubMed PMID: 22232059. Pubmed Central PMCID: PMC3490385.

43. Zhuang Y, Peng H, Mastej V, Chen W. MicroRNA Regulation of Endothelial Junction Proteins and Clinical Consequence. Mediators Inflamm. 2016;2016:5078627. PubMed PMID: 27999452. Pubmed Central PMCID: PMC5143735.

44. Lukamowicz-Rajska M, Mittmann C, Prummer M, Zhong Q, Bedke J, Hennenlotter J, et al. MiR-99b-5p expression and response to tyrosine kinase inhibitor treatment in clear cell renal cell carcinoma patients. Oncotarget. 2016 Nov 29;7(48):78433–47. PubMed PMID: 27738339.

45. Jin Y, Tymen SD, Chen D, Fang ZJ, Zhao Y, Dragas D, et al. MicroRNA-99 family targets AKT/mTOR signaling pathway in dermal wound healing. PLoS One. 2013;8(5):e64434. PubMed PMID: 23724047. Pubmed Central PMCID: PMC3665798.

46. Schweighofer B, Testori J, Sturtzel C, Sattler S, Mayer H, Wagner O, et al. The VEGF-induced transcriptional response comprises gene clusters at the crossroad of angiogenesis and inflammation. Thromb Haemost. 2009 Sep;102(3):544–54. PubMed PMID: 19718476. Pubmed Central PMCID: PMC2886966.

47. Minamino T, Miyauchi H, Yoshida T, Ishida Y, Yoshida H, Komuro I. Endothelial Cell Senescence in Human Atherosclerosis. Role of Telomere in Endothelial Dysfunction. 2002;105(13):1541–4.

48. Kurz DJ, Decary S, Hong Y, Trivier E, Akhmedov A, Erusalimsky JD. Chronic oxidative stress compromises telomere integrity and accelerates the onset of senescence in human endothelial cells. Journal of Cell Science. 2004;117(11):2417–26.

49. Dzikiewicz-Krawczyk A. The importance of making ends meet: mutations in genes and altered expression of proteins of the MRN complex and cancer. Mutat Res. 2008 Sep-Oct;659(3):262–73. PubMed PMID: 18606567.

50. O’Malley BW, Jr., Li D, Carney J, Rhee J, Suntharalingam M. Molecular disruption of the MRN(95) complex induces radiation sensitivity in head and neck cancer. Laryngoscope. 2003 Sep;113(9):1588–94. PubMed PMID: 12972939.

51. Araki K, Yamashita T, Reddy N, Wang H, Abuzeid WM, Khan K, et al. Molecular disruption of NBS1 with targeted gene delivery enhances chemosensitisation in head and neck cancer. Br J Cancer. 2010 12/07/print;103(12):1822–30.

52. Mun GI, Boo YC. Identification of CD44 as a senescence-induced cell adhesion gene responsible for the enhanced monocyte recruitment to senescent endothelial cells. American Journal of Physiology - Heart and Circulatory Physiology. 2010;298(6):H2102–H11.

53. Flynn KM, Michaud M, Canosa S, Madri JA. CD44 regulates vascular endothelial barrier integrity via a PECAM-1 dependent mechanism. Angiogenesis. 2013 Jul;16(3):689–705. PubMed PMID: 23504212.

54. Orian-Rousseau V, Chen L, Sleeman JP, Herrlich P, Ponta H. CD44 is required for two consecutive steps in HGF/c-Met signaling. Genes & Development. 2002 December 1, 2002;16(23):3074–86.

55. DeLisser HM. CD44: target for antiangiogenesis therapy. Blood. 2009;114(25):5114–5.

56. Li Y, Shen Y, Hohensinner P, Ju J, Wen Z, Goodman Stuart B, et al. Deficient Activity of the Nuclease MRE11A Induces T Cell Aging and Promotes Arthritogenic Effector Functions in Patients with Rheumatoid Arthritis. Immunity. 2016 10/18/;45(4):903–16.

57. Rai R, Hu C, Broton C, Chen Y, Lei M, Chang S. NBS1 Phosphorylation Status Dictates Repair Choice of Dysfunctional Telomeres. Molecular Cell. 2017 3/2/;65(5):801–17.e4.

58. Wu Y, Xiao S, Zhu X-D. MRE11-RAD50-NBS1 and ATM function as co-mediators of TRF1 in telomere length control. Nat Struct Mol Biol. 2007 09//print;14(9):832–40.

59. Wilson R, Espinosa-Diez C, Kanner N, Chatterjee N, Ruhl R, Hipfinger C, et al. MicroRNA regulation of endothelial TREX1 reprograms the tumour microenvironment. Nature Communications. 2016 11/25 06/27/received 10/18/accepted;7:13597. PubMed PMID: PMC5133658.

60. Anand S, Majeti BK, Acevedo LM, Murphy EA, Mukthavaram R, Scheppke L, et al. MicroRNA-132-mediated loss of p120RasGAP activates the endothelium to facilitate pathological angiogenesis. Nature medicine. 2010 Aug;16(8):909–14. PubMed PMID: 20676106. Pubmed Central PMCID: 3094020.

61. Kelley KA, Ruhl RA, Rana SR, Dewey E, Espinosa C, Thomas CRJ, et al. Understanding and Resetting Radiation Sensitivity in Rectal Cancer. Annals of Surgery. 2017;266(4):610–6. PubMed PMID: 00000658-201710000-00008.

62. Shiao SL, Ruffell B, DeNardo DG, Faddegon BA, Park CC, Coussens LM. TH2-Polarized CD4(+) T Cells and Macrophages Limit Efficacy of Radiotherapy. Cancer immunology research. 2015 May;3(5):518–25. PubMed PMID: 25716473. Pubmed Central PMCID: 4420686.

